# Phytocystatin 6 is a context-dependent, tight-binding inhibitor of *Arabidopsis thaliana* legumain isoform β

**DOI:** 10.1101/2023.05.22.541692

**Authors:** Naiá P. Santos, Wai T. Soh, Fatih Demir, Raimund Tenhaken, Peter Briza, Pitter F. Huesgen, Hans Brandstetter, Elfriede Dall

**Affiliations:** Department of Biosciences and Medical Biology, University of Salzburg, 5020 Salzburg, Austria; Central Institute for Engineering, Electronics and Analytics, ZEA-3, Forschungszentrum Jülich, 52428 Jülich, Germany; Department of Environment and Biodiversity, University of Salzburg, 5020 Salzburg, Austria; CECAD, Medical Faculty and University Hospital, University of Cologne, 50931 Cologne, Germany; Institute for Biochemistry, Faculty of Mathematics and Natural Sciences, University of Cologne, 50674 Cologne, Germany; Max Planck Institute for Multidisciplinary Sciences, D-37077 Göttingen, Germany; Department of Biomedicine, Aarhus University, 8000 Aarhus C, Denmark

## Abstract

Plant legumains are crucial for processing seed storage proteins and are critical regulators of plant programmed cell death. Although research on legumains boosted recently, little is known about their activity regulation. In our study, we used pull-down experiments to identify AtCYT6 as a natural inhibitor of legumain isoform β (AtLEGβ) in *Arabidopsis thaliana*. Biochemical analysis revealed that AtCYT6 inhibits both AtLEGβ and papain-like cysteine proteases through two cystatin domains. The N-terminal domain inhibits papain-like proteases, while the C-terminal domain inhibits AtLEGβ. Furthermore, we showed that AtCYT6 interacts with legumain in a substrate-like manner, facilitated by a conserved asparagine residue in its reactive center loop. Complex formation was additionally stabilized by charged exosite interactions, contributing to pH-dependent inhibition. Processing of AtCYT6 by AtLEGβ suggests a context-specific regulatory mechanism with implications for plant physiology, development, and programmed cell death. These findings enhance our understanding of AtLEGβ regulation and its broader physiological significance.

## INTRODUCTION

*Arabidopsis thaliana* expresses four isoforms of the cysteine protease legumain (C13 family, EC 3.4.22.34) denoted as AtLEGα, -β, -γ and -δ. Due to their organ-specific roles, they are assorted as vegetative-(AtLEGγ and AtLEGα) and seed-type legumains (AtLEGβ and AtLEGδ) (1,2). Together they are accountable for a myriad of defense and developmental responses such as the physiological and stress-induced programmed cell death (PCD), seed papain-like cysteine proteases (PLCP, clan C1) storage mobilization and processing of vacuolar proteins, the latter leading to their synonymous naming as vacuolar processing enzymes (VPEs) (3–10). AtLEGβ is the most relevant isoform for seed storage processing and mobilization. Naturally occurring *A. thaliana* specimens with dysfunctional *atlegβ* genes showed aberrant seed storage processing (11). Furthermore, AtLEGβ-null mutants display impaired pollen fertility and tapetal cell degradation (12). AtLEGβ is synthesized as an inactive proenzyme (proAtLEGβ) consisting of a caspase-like catalytic domain and a death-domain-like C-terminal prodomain. Activation proceeds via the autocatalytic, pH-dependent removal of the prodomain. Activated AtLEGβ specifically cleaves after asparagine, and to a lesser extent aspartate, residues. It is stable and proteolytically active at acidic pH, displays transpeptidase and ligase activity at near-neutral pHs; over time, it undergoes irreversible denaturation at pH > 6.0 (13–15). Given its versatile activities, tight regulation is required and may be conferred *in vivo* by cystatins.

Cystatins are reversible competitive inhibitors of PLCPs, which are well characterized in mammals but less so in plants. Human cystatins are subdivided into type I, II and III cystatins. Type I cystatins are intracellular, un-glycosylated small proteins lacking disulfide bonds, in contrast to type II cystatins which are secreted, typically glycosylated and disulfide-linked. Single-domain type II cystatins harbor a legumain inhibitory site, distinct from the PLCP site. Type III cystatins are multidomain cystatins with or without inhibitory activity encoded within their individual domains (16,17). Phytocystatins (phycys), the cystatins from plants, can be classified according to similar criteria as in mammals, except for type II phycys. Type II phycys are two-domain inhibitors that target PLCPs and legumains via two distinct cystatin-like domains (18). While the N-terminal domain encodes PLCP inhibition, legumain inhibition is encoded in the C-terminal domain. The cystatin superfamily shares a highly conserved fold consisting of a five stranded antiparallel β-sheet wrapped around a central α-helix (19). Cystatins bind to PLCPs in the so-called elephant-trunk binding mode, where the strictly conserved and critical inhibition determinants consist of an N-terminal glycine, the Q-X-V-X-G motif on the L1-loop, and the P-W motif on the L3 loop (19,20). By contrast, the binding mode of legumain and type II cystatins has been reported by Dall *et al*. as a substrate-like interaction between human cystatin E (hCE) and human legumain (hLEG) where the P1-Asn on the reactive center loop (RCL) of hCE inserts into the S1 pocket of hLEG, thus blocking the access of the substrate. The interaction with the enzyme is strengthened by the legumain exosite loop (LEL), which in hCE is long and predominantly hydrophobic (21). Furthermore, complex formation is tightly regulated by a pH-dependent equilibrium of RCL cleavage at acidic pH and re-ligation at near neutral pH. In spite of a relatively broad knowledge of the interaction of hLEG and human cystatins, there is little data available on the interplay of plant legumains and their endogenous regulators, e.g. phytocystatins. We hypothesized that *A. thaliana* phytocystatins regulate AtLEGβ activity and were able to characterize the type II phytocystatin 6 (AtCYT6) as a potent substrate-like inhibitor of the enzyme. The finding presented here on the interaction of AtCYT6 with AtLEGβ contribute to our understanding of AtLEGβ activity *in vivo*, and to the identification of structural determinants of type II phytocystatins important for inhibition.

## RESULTS

### Phytocystatin 6 is a candidate inhibitor of *Arabidopsis thaliana* legumain β (AtLEGβ)

To study how the activity of AtLEGβ is regulated in *Arabidopsis thaliana* we were looking out for potential inhibitors of its enzymatic activities. In mammals, legumain activity is inhibited by the endogenous family II cystatins, including cystatin C, M/E, and F (22). Similarly, type II phycys were shown in previous studies to inhibit legumain from rice (23), barley (24), and humans (18). Till now, however, not much is known on phycys in *A. thaliana*. In 2005, Martinez and colleagues identified seven phytocystatin candidate genes in *A. thaliana* (AtCYT1-7) through a comparative phylogenetic analysis (25). Four of them, AtCYT1-3 and AtCYT6, have in the meantime been confirmed on protein level (26–30). To find out if and which of *A. thaliana* phytocystatin candidate genes harbour the characteristic tandem two domain architecture, we used AlphaFold to predict their protein structures (31,32). Intriguingly, although AtCYT6 and AtCYT7 were expected to harbor a two-domain architecture based on their primary sequence (Fig. S1), only AtCYT6 resulted in an AlphaFold model that contained two folded domains with cystatin-like architecture (Fig. 1A). AtCYT7, as well as AtCYT1-5, contained only one fully folded cystatin domain at the N-terminal end. In the model, the characteristic carboxy-terminal extension of type II phycys, although present in the sequence of AtCYT7, was primarily constituted of disordered regions (not shown). Overall, the AtCYT6 AlphaFold model had a good per-residue confidence score, with 70% of all residues having a predicted local distance difference test (pLDDT) value higher than 70. This translated in *confident* (90 > pLDDT > 70) or *very high confident* predictions (pLDDT > 90). The exceptions were the N-terminal region up to residue Gly39 (including the signal peptide) and the interdomain linker (Ala128-Glu144) connecting the two predicted cystatin domains. The individual N- and C-terminal domains (NTD and CTD) both displayed an α-helix wrapped by an anti-parallel β-sheet, the characteristic tertiary structure of members of the cystatin family (19). The superposition of modeled NTD and CTD yielded a Cα backbone RMSD of 2.20 Å (Fig. 1B). Interestingly, superposition and sequence alignments indicated a potential legumain reactive center loop (RCL) harboring an asparagine as putative P1-residue, as seen in the legumain inhibitory human cystatin M/E and C, in both the NTD and the CTD (Fig. 1B, C) (21). Furthermore, sequence analysis also uncovered a L/VGG motif on the NTD, which was previously linked to PLCP inhibition in other cystatins. Based on these findings, we hypothesized that AtCYT6 is a type II phytocystatin capable of inhibiting both *A. thaliana* legumains and PLCPs.

**Figure 1:**
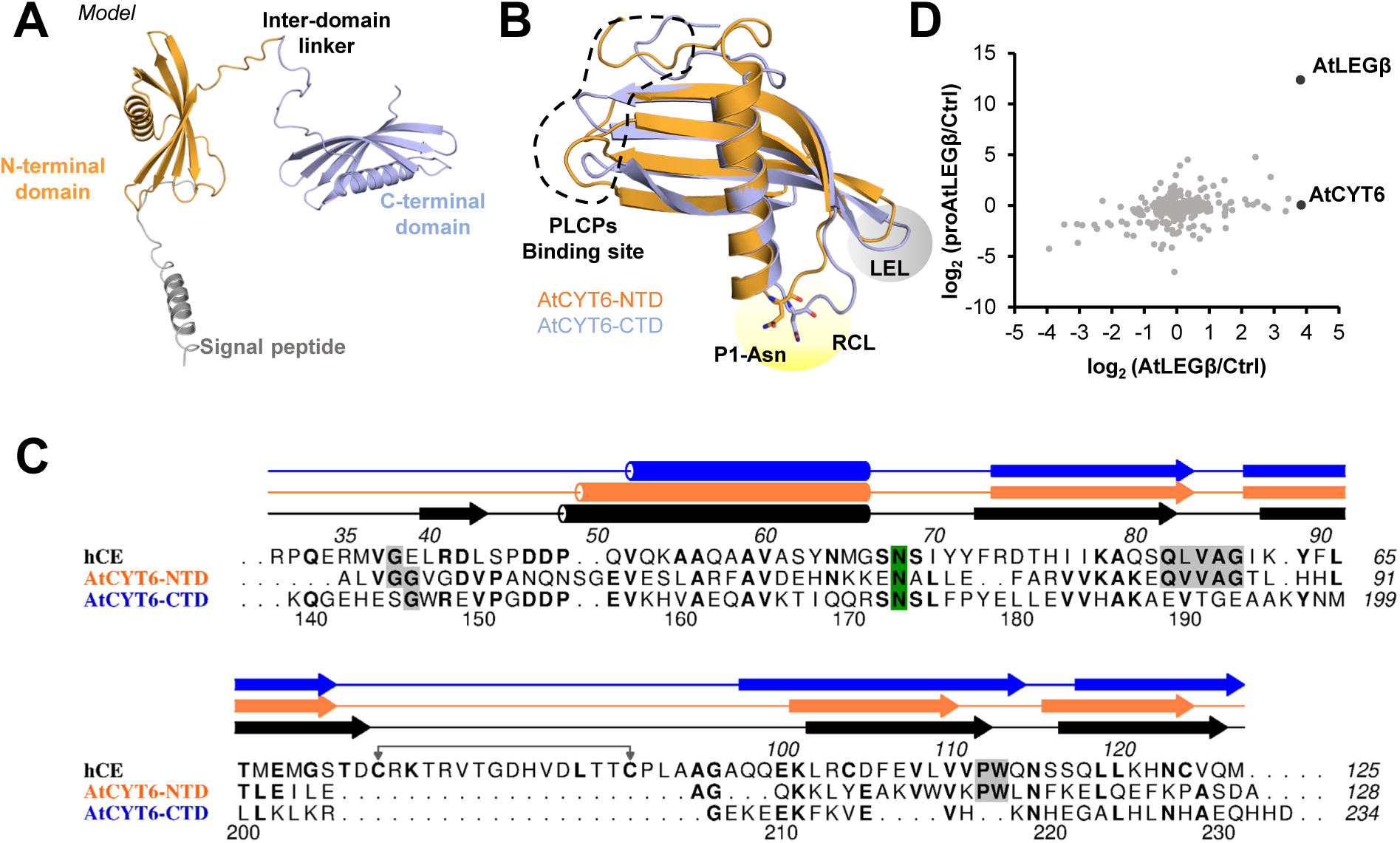
AtCYT6 is a two-domain phytocystatin that interacts with AtLEGβ. **(A)** AlphaFold model of full-length AtCYT6 depicting the N-terminal signal peptide (grey), the N-terminal domain (orange), the inter-domain linker and the C-terminal domain (blue). **(B)** Structure alignment of the AlphaFold models for the N- (orange, AtCYT6-NTD) and C-terminal (blue, AtCYT6-CTD) domains. P1-Asn residues are shown in sticks; reactive center loops (RCL) are highlighted in yellow; legumain exosite loops are highlighted in grey; the binding site for PLCPs in the N-terminal domain is indicated by dashed lines. **(C)** Sequence alignment of hCE (black), AtCYT6-NTD (orange) and AtCYT6-CTD (blue) generated with Aline. Secondary structure elements are indicated, the P1-Asn residues are highlighted in green, and PLCPs inhibition motifs in grey. **(D)** Correlation of pull-down data of AtLEGβ/beads and proAtLEGβ/beads. Extracts of *A. thaliana* leaves of a VPE_0_ strain were spiked with recombinant His_6_-tagged (pro)AtLEGβ and co-precipitated via Ni^2+^-affinity chromatography. AtCYT6 and AtLEGβ-derived peptides displayed the highest log_2_-increase relative to control beads.

### CYT6 is an interaction partner of AtLEGβ

To find out whether AtCYT6 was indeed an inhibitor of AtLEGβ *in planta*, pull-down assays were performed. For that purpose, extracts of *A. thaliana* leaves of a VPE_0_ strain lacking the expression of all four legumain isoforms were used. The extracts were spiked with recombinant inactive proAtLEGβ or active AtLEGβ, both harboring an N-terminal His_6_-tag, which was used to recover the proteins via magnetic Ni^2+^-beads. In control experiments the extracts were spiked with buffer instead of AtLEGβ. Differential triplex labeling of the proAtLEGβ, AtLEGβ, and buffer control samples followed by mass spectrometry analysis confirmed that AtLEGβ was significantly increased in both protease-treated samples relative to the buffer control (Fig. 1D). Importantly, AtCYT6 showed the highest log2-fold change in the sample spiked with AtLEGβ but was not co-purified with proAtLEGβ, suggesting that the binding of AtCYT6 to AtLEGβ requires an interaction surface that is available in AtLEGβ but not in proAtLEGβ, e.g., the active site. This finding corroborated AtCYT6 as an interaction partner and likely physiologic inhibitor of AtLEGβ.

### The CTD of AtCYT6 inhibits AtLEGβ

To further confirm that AtCYT6 was indeed binding to and inhibiting AtLEGβ, we recombinantly expressed and purified to homogeneity the full-length AtCYT6 lacking the N-terminal signal peptide (A35-D235, AtCYT6-FL), the N-terminal domain alone (A35-D127, AtCYT6-NTD) and the C-terminal domain alone (E142-D235, AtCYT6-CTD). The identity of the recombinant proteins was confirmed by mass spectrometry (Table S1). As a first assessment of legumain interaction sites, we carried out co-migration assays. Specifically, we co-incubated active AtLEGβ with AtCYT6-FL, AtCYT6-NTD or AtCYT6-CTD separately and subjected the samples to size exclusion chromatography (SEC) experiments. The AtCYT6-FL co-migrated with AtLEGβ, reiterating its binding properties to AtLEGβ seen in the pull-down assay (Fig. 2A). When we tested the AtCYT6-NTD and AtCYT6-CTD individually, co-migration with the enzyme was only observed with the AtCYT6-CTD (Fig. 2B,C). Subsequently, we tested whether the AtCYT6 constructs were indeed inhibitors of legumain activity. To that end, we co-incubated active AtLEGβ with the individual constructs and monitored the turnover of the legumain-specific AAN-AMC substrate. In line with our SEC experiments, we found that AtCYT6-FL and AtCYT6-CTD inhibited the proteolytic activity of AtLEGβ, but not the AtCYT6-NTD (Fig. 3A). In a next step, we compared the affinities of AtCYT6-FL and AtCYT6-CTD by determining their inhibition constants (K_i_). The constructs displayed similar, sub-nanomolar affinities towards AtLEGβ (K_i AtCYT6-FL_ = 0.29 ± 0.05 nM; K_i AtCYT6-CTD_ = 0.61 ± 0.23 nM) (Table 1 and Fig. S2A,B). Withal, a second variant of the C-terminal domain (A128-D235, AtCYT6-CTD_long_) carrying the interdomain linker was produced and inhibited the enzyme with similar K_i_ as AtCYT6-FL and AtCYT6-CTD (K_i AtCYT6-CTDlong_ = 0.68 ± 0.20 nM). Based on these results we concluded that AtCYT6 was a high affinity physiologic inhibitor of AtLEGβ, and that inhibition was mediated by its C-terminal domain.

**Figure 2:**
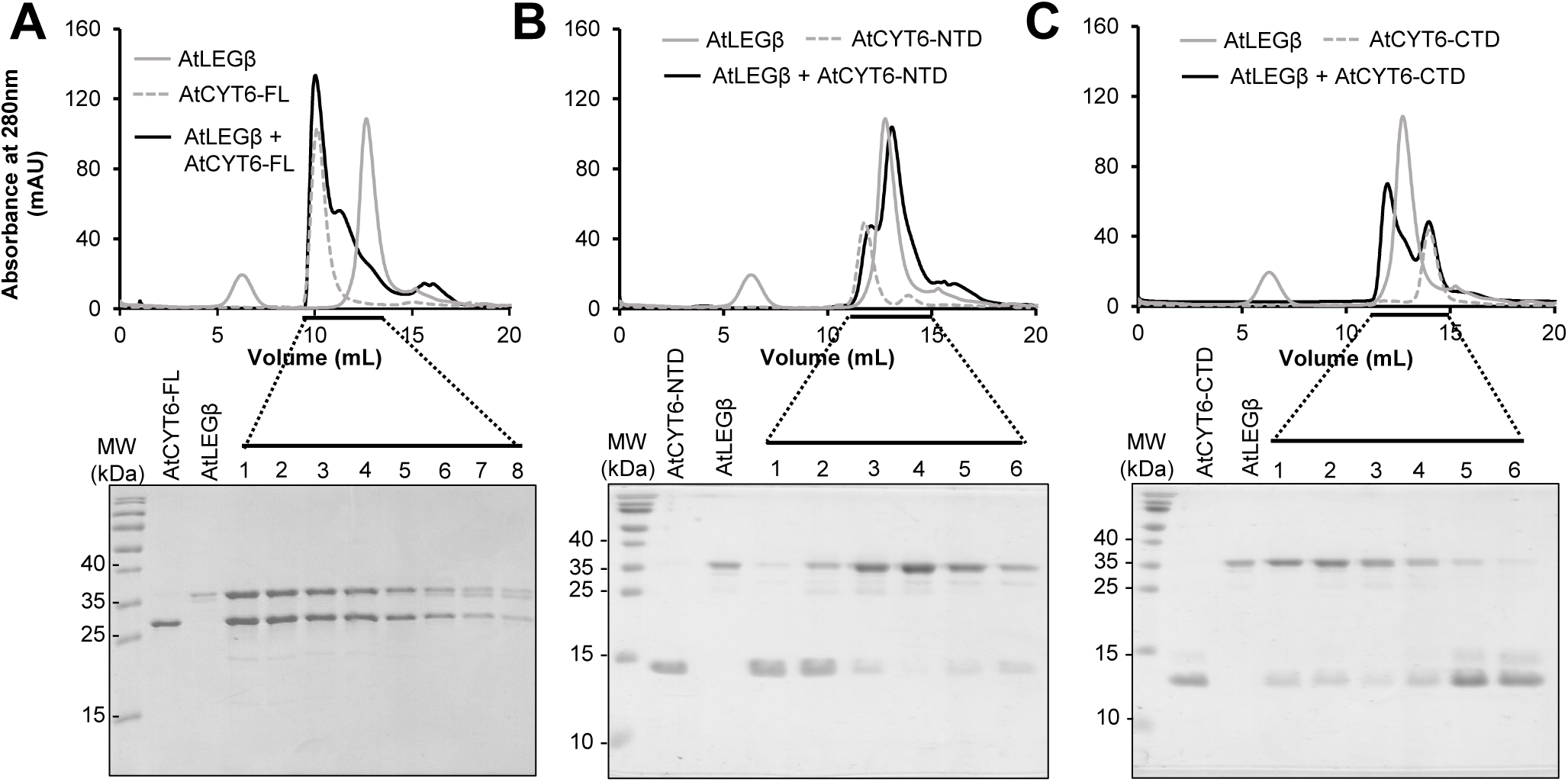
The interaction of AtCYT6 with AtLEGβ is mediated by the C-terminal domain (AtCYT6-CTD). AtLEGβ was inhibited with methyl-methanethiosulfonate (MMTS), incubated with **(A)** alkylated AtCYT6-FL, **(B)** AtCYT6-NTD or **(C)** AtCYT6-CTD at pH 5.5 and complex formation was analyzed by size exclusion chromatography. Indicated fractions were analyzed by SDS-PAGE. Control reactions contained only AtLEGβ or the respective AtCYT6 construct.

**Figure 3:**
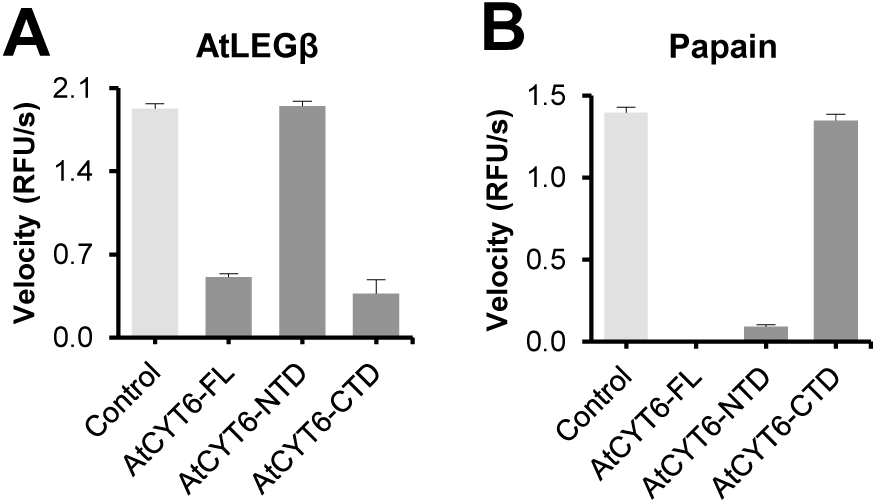
AtCYT6 is a potent inhibitor of AtLEGβ and papain. **(A)** The turnover of the AAN-AMC substrate by AtLEGβ was assayed at pH 5.5 in the absence (light grey) and presence (dark grey) of AtCYT6-FL, AtCYT6-NTD or AtCYT6-CTD. **(B)** Turnover of the FR-AMC substrate by papain measured at pH 6.5.

**Table 1:**
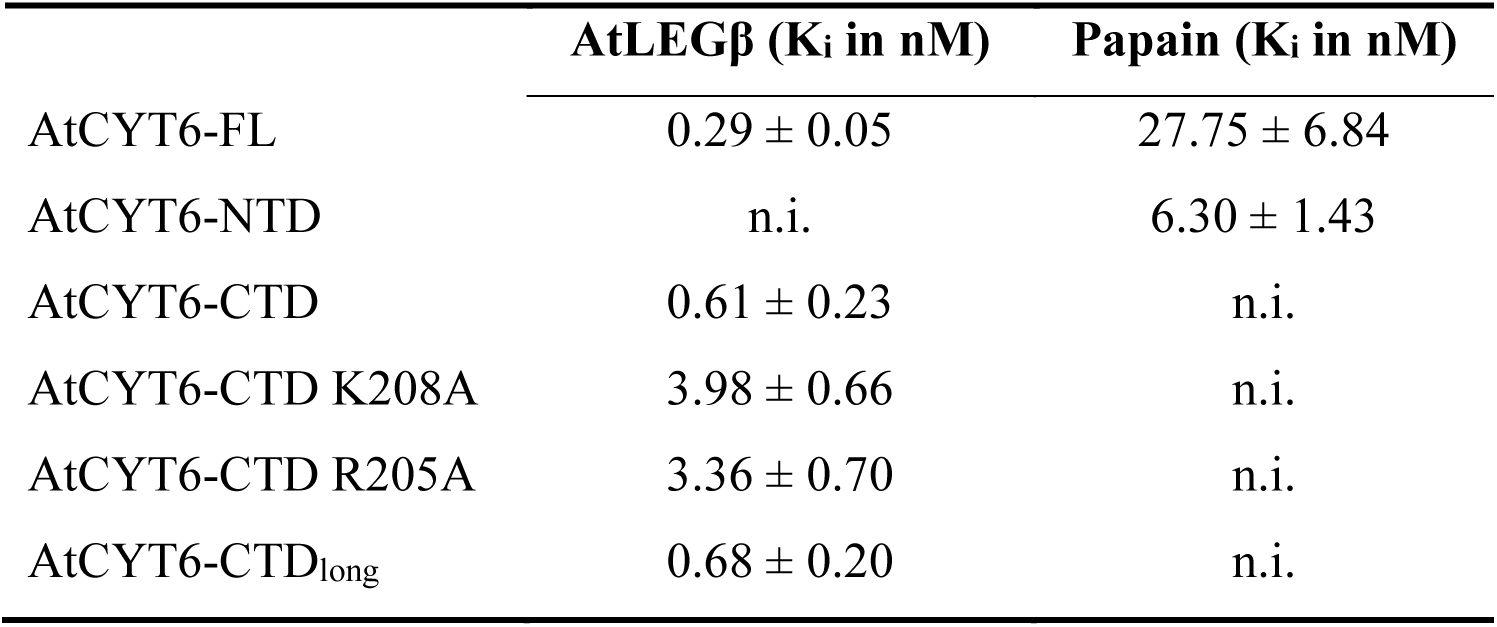
K_i_ values of different AtCYT6 constructs against AtLEGβ and papain.

| | AtLEGβ ($K_i$ in nM) | Papain ($K_i$ in nM) |
| --- | --- | --- |
| AtCYT6-FL | $0.29 \pm 0.05$ | $27.75 \pm 6.84$ |
| AtCYT6-NTD | n.i. | $6.30 \pm 1.43$ |
| AtCYT6-CTD | $0.61 \pm 0.23$ | n.i. |
| AtCYT6-CTD K208A | $3.98 \pm 0.66$ | n.i. |
| AtCYT6-CTD R205A | $3.36 \pm 0.70$ | n.i. |
| AtCYT6-CTD<sub>long</sub> | $0.68 \pm 0.20$ | n.i. |

### AtCYT6-NTD inhibits papain-like cysteine proteases (PLCPs)

Since cystatins are well established inhibitors of PLCPs, in a next step we tested if our recombinant AtCYT6 constructs would also be PLCP inhibitors. The L/VGG motif at the N-terminus of inhibitory cystatins is critical for inhibition of PLCPs by other cystatins (33). The absence of such a motif in the AtCYT6-CTD and its presence at the AtCYT6-NTD suggested that the NTD harbored a PLCP-inhibitory activity (Fig. 1C). To confirm this hypothesis, papain was used as a model for the PLCP family. The turnover of its FR-AMC substrate was monitored upon incubation of papain with AtCYT6-FL, AtCYT6-NTD or AtCYT6-CTD. Indeed, AtCYT6-FL and AtCYT6-NTD exhibited high binding affinities towards papain (K_i AtCYT6-FL_ = 27.75 ± 6.84 nM; K_i AtCYT6-NTD_ = 6.30 ± 1.43 nM), but not AtCYT6-CTD (Fig. 3B, Table 1 and Fig. S2C,D). In accordance with our hypotheses, the N- and C-terminal domains of AtCYT6 exhibited distinct inhibitory properties towards specific cysteine proteases.

### AtLEGβ selectively binds to AtCYT6-CTD *in vivo*

To further validate the selectivity of AtCYT6-CTD as legumain inhibitory domain *in vivo*, we performed co-immunoprecipitation assays (CoIPs) using *A. thaliana* seed extracts and polyclonal rabbit antibodies raised against AtCYT6-NTD or AtCYT6-CTD. To precipitate endogenous AtCYT6, we immobilized anti-AtCYT6-NTD or anti-AtCYT6-CTD antibodies on protein G-coupled sepharose beads. Subsequently we loaded the soluble fractions of seed extracts prepared at pH 5.5, followed by washing and elution of captured proteins. Control samples contained protein G-coupled sepharose beads only. After digestion of the samples with trypsin, they were analyzed by label-free mass spectrometry (Fig. 4A). The abundance of proteins identified in each sample was calculated as log_2_ ratios between label-free quantification intensities of test samples and control beads. Importantly, both antibodies, anti-AtCYT6-NTD and anti-AtCYT6-CTD, yielded cystatin 6-derived peptides as the most abundant species (positive control). Interestingly, only anti-AtCYT6-CTD but not anti-AtCYT6-NTD antibodies co-precipitated endogenous AtLEGβ with endogenous AtCYT6. This experiment confirmed that AtCYT6 is a physiologic inhibitor of AtLEGβ in *A. thaliana* seeds and that also *in vivo* their interaction is mediated by the AtCYT6-CTD. Furthermore, we suggest that AtCYT6 was at least partially processed in our CoIP samples, e.g. in the inter-domain linker and that therefore the anti-AtCYT6-NTD antibodies were not able to co-elute AtLEGβ. This suggestion was supported by western blot experiments of the elutions of the CoIPs, which showed no or little protein at the expected height of AtCYT6-FL (Fig. S3). Instead, we observed bands corresponding in size to AtCYT6-NTD or AtCYT6-CTD, respectively. Importantly, no other cystatin isoform besides cystatin 6 was co-purified in the CoIPs, confirming the specificity of the antibodies used. To further confirm the observed *in vivo* selectivity, we repeated the CoIP experiments using beads where recombinant AtCYT6-NTD or AtCYT6-CTD were loaded to the immobilized anti-AtCYT6-NTD or anti-AtCYT6-CTD antibodies respectively. Importantly, AtLEGβ was recovered at 20-times higher intensity using anti-AtCYT6-CTD loaded with recombinant AtCYT6-CTD than with anti-AtCYT6-NTD loaded with recombinant AtCYT6-NTD (Fig. 4B). Further analysis of our CoIP experiments additionally uncovered specific interaction partners of AtCYT6-NTD, like the PLCPs SAG12, RD19D and a cysteine proteinase superfamily protein (Q9LNC1) with so far unknown function (Fig. 4). All of them were specifically observed when the anti-AtCYT6-NTD antibody was used as a bait, with or without AtCYT6-NTD. This suggested that AtCYT6-NTD specifically inhibits PLCPs also *in vivo* and implies that AtCYT6 may be a physiological regulator of SAG12 and RD19D activities.

**Figure 4:**
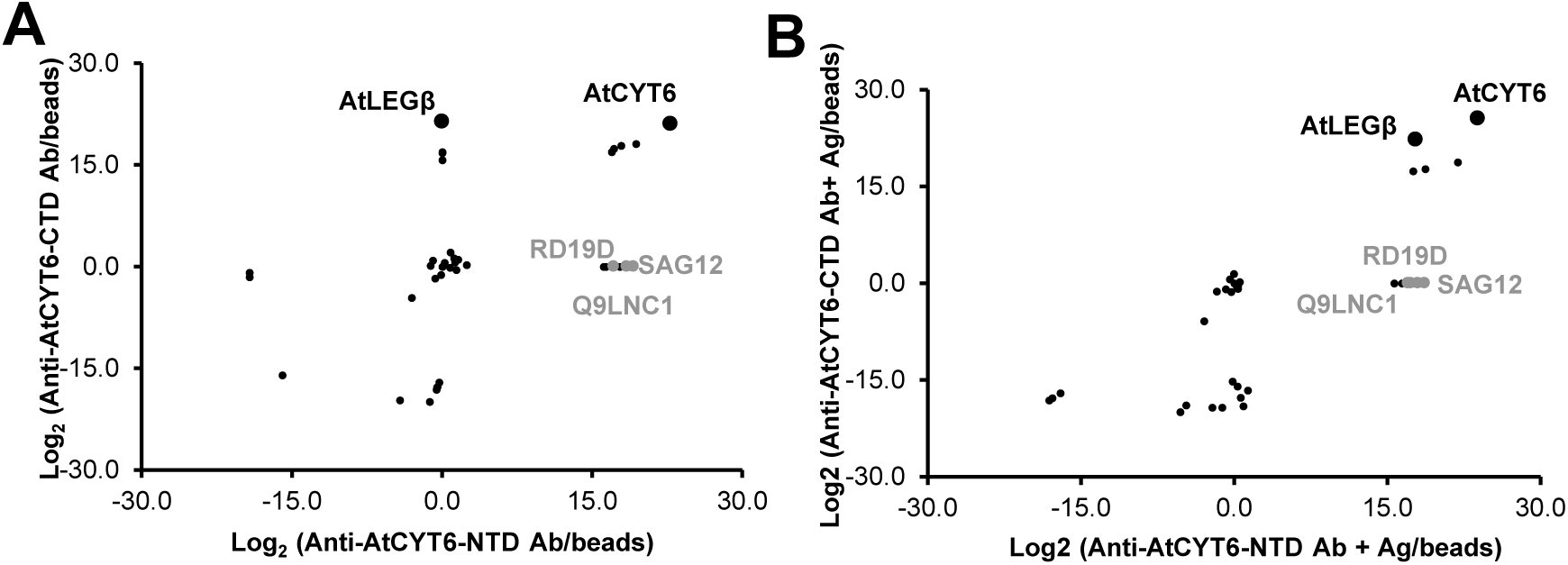
Co-immunoprecipitation assays cross-confirmed interaction of AtLEGβ and AtCYT6 in vivo. **(A)** Polyclonal anti-AtCYT6-NTD (Anti-AtCYT6-NTD) or anti-AtCYT6-CTD antibodies were cross-linked to protein G-beads and incubated with seed extract at pH 5.5. Protein abundances are shown as log_2_ of the ratio of label-free quantification intensities of test samples and control beads (null values were substituted by 1 to enable the calculations). **(B)** Same as **(A)** but antibodies were loaded with the respective recombinantly expressed antigen before incubation with the seed extract. AtCYT6-derived peptides were the most abundant in both experiments. AtLEGβ was co-eluted in the presence of the anti-AtCYT6-CTD antibody and with antibodies loaded with recombinant AtCYT6-NTD or AtCYT6-CTD.

### AtCYT6 is a substrate-like inhibitor of AtLEGβ

Knowing that the C-terminal domain of AtCYT6 was the actual legumain inhibitor, we were in a next step wondering whether it would exploit a substrate-like mode of interaction, as seen in human legumain-inhibitory cystatins (21). Along that line, based on our model, Asn173 could serve as a P1-residue. To test this hypothesis, we prepared an AtCYT6-CTD-N173A mutant and tested whether it would be inhibiting AtLEGβ. Using AAN-AMC as a substrate, we indeed found that the mutant lost its ability to inhibit AtLEGβ activity (Fig. 5A). This finding confirmed the essential role of Asn173 within the proposed RCL as a mediator of the interaction of AtCYT6-CTD with AtLEGβ. Since an asparagine was indeed essential for the interaction, we then hypothesized that Asn173 serves as a P1 residue binding in a productive, substrate-like orientation to the S1-pocket on AtLEGβ. To test this hypothesis, we co-incubated AtLEGβ with AtCYT6-FL or the AtCYT6-CTD at pH 5.5 in a 1:5 molar ratio (enzyme:inhibitor) and analyzed the reaction for cleavage products using SDS-PAGE and mass spectrometry. Indeed, we found that both AtCYT6 variants were processed by AtLEGβ (Fig. 5B). Interestingly, cleavage was only observed for less than 20 % of the AtCYT6-FL or AtCYT6-CTD proteins, leaving 80 % of these proteins intact. This led us to the conclusion that AtCYT6 remained in a stable complex with AtLEGβ even though partial AtCYT6 cleavage by AtLEGβ was observed. Additional mass spectrometry experiments confirmed that Asn173 was the most prominent cleavage site in both constructs (Table S1 and Fig. 5C).

**Figure 5:**
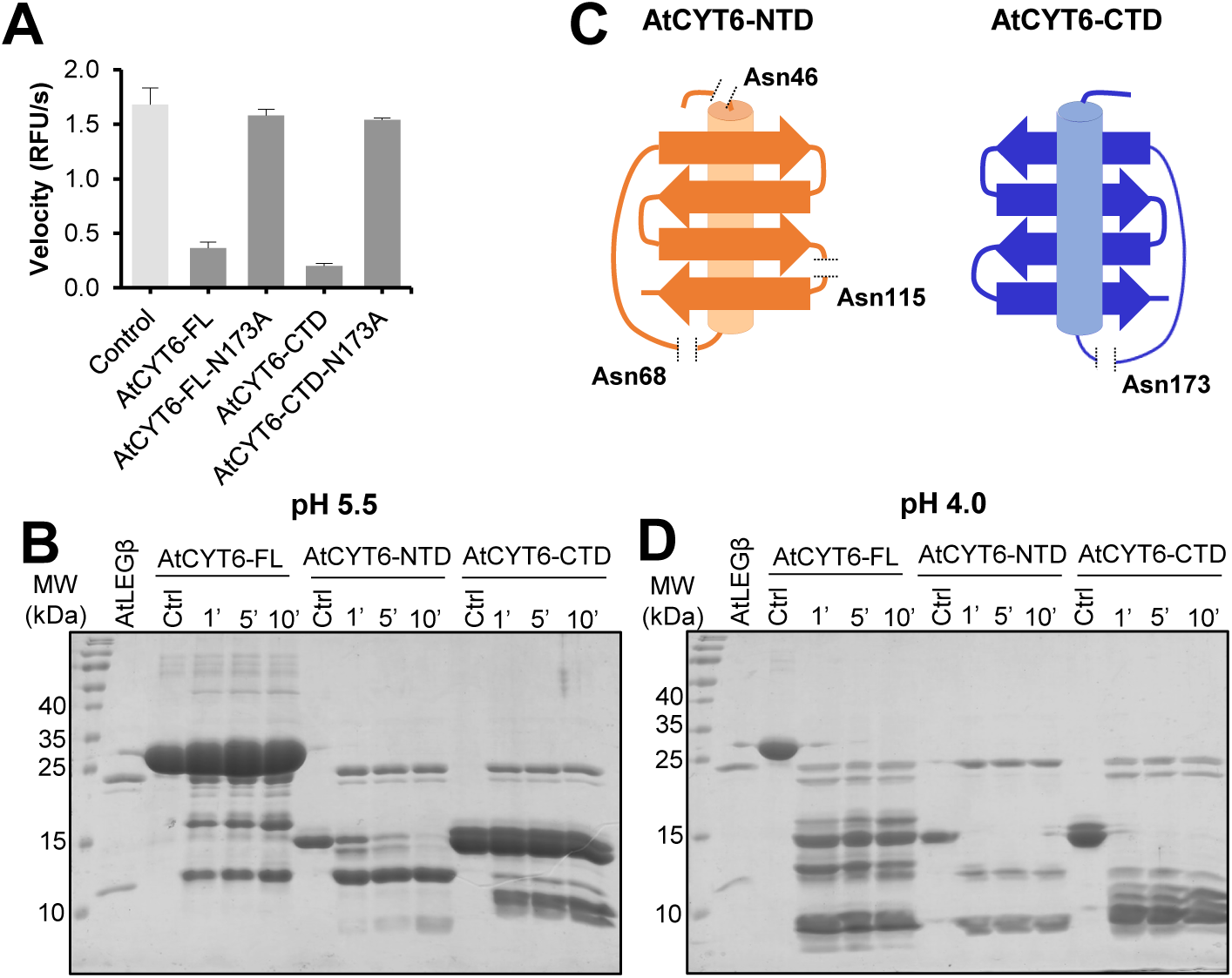
The interaction of AtCYT6 with AtLEGβ is mediated by P1-Asn173. **(A)** Reaction velocities for the turnover of the AAN-AMC substrate by AtLEGβ in the absence (light grey) and presence (dark grey) of AtCYT6-FL, AtCYT6-FL-N173A, AtCYT6-CTD or AtCYT6-CTD-N173A. **(B)** SDS-PAGE of AtCYT6-FL, AtCYT6-NTD or AtCYT6-CTD after incubation with AtLEGβ in a 1:5 molar ratio at pH 5.5 after 1, 5 and 10 minutes of incubation. **(C)** Schematic representation of the cleavage sites of AtLEGβ within AtCYT6-NTD (orange) and AtCYT6-CTD (blue) identified by mass spectrometry. **(D)** Same as **(B)** but the incubation was done at pH 4.0.

Although AtCYT6-NTD harbored an asparagine residue (Asn68) at a position structurally equivalent to Asn173 on AtCYT6-CTD, it did not show inhibition of AtLEGβ. To test whether AtCYT6-NTD was a substrate to AtLEGβ rather than an inhibitor, we co-incubated them at pH 5.5 and similarly analyzed the reaction by SDS-PAGE and mass spectrometry. Interestingly we found that in contrast to the other AtCYT6 constructs, the N-terminal domain was processed completely within less than 10 min of incubation (Fig. 5B). Mass spectrometry experiments confirmed that cleavage occurred after the P1-Asn68 residue (Table S1). Additional cleavage events resulted in complete fragmentation of the N-terminal domain by AtLEGβ (Fig. 5C). These experiments confirmed our hypothesis that AtCYT6-NTD was rather a substrate to AtLEGβ than an inhibitor.

### The double basic Arg205-Lys208 motif establishes pH-dependent exosite interactions

Interestingly, both cystatin-like domains on AtCYT6 harbor an asparagine on the RCL that may be recognized like a substrate by AtLEGβ. However, while the N-terminal domain is a substrate, the C-terminal domain additionally functions as an inhibitor. Prompted by this observation, we wondered which feature was turning the C-terminal domain into an inhibitor. To address this question, we prepared models of AtLEGβ in complex with AtCYT6-FL, AtCYT6-NTD or AtCYT6-CTD using AlphaFold-Multimer and compared them to the crystal structure of hLEG in complex with hCE (PDB 4N6O, Fig. 6A and S3) (32). Importantly, the modeled structure of AtLEGβ was in good agreement with a crystallographic structure of proAtLEGβ (PDB 6YSA) which we previously determined (Cα RMSD = 0.38 Å; determined with PyMOL). Also, the Cα RMSD between the C-terminal domain modeled in complex with AtLEGβ in presence and absence of the N-terminal domain, i.e. AtCYT6-FL and AtCYT6-CTD, was < 0.1 Å (Fig. S4A), which is in accordance with our experimental data supporting similar K_i_ values for both constructs against AtLEGβ. The models suggested that the RCL interaction was similar in hCE and AtCYT6-CTD. The RCL of AtCYT6-CTD established substrate-like interactions to the S3-S2‘-substrate binding sites on AtLEGβ, where Asn173 serves as a P1 residue (Fig. 6A,B, S4B). Interestingly, AlphaFold-Multimer positioned AtCYT6-NTD in an ‘upside-down’ orientation onto the active site of AtLEGβ, with Asn48 positioned to the S1-pocket (Fig. S4C). These models are in qualitative agreement with our experimental data, which showed that only AtCYT6-CTD is an effective inhibitor of AtLEGβ, but not AtCYT6-NTD. To force AlphaFold-Multimer to position Asn68 of AtCYT6-NTD to the active site of AtLEGβ we replaced all asparagine residues that were identified as cleavage sites by mass spectrometry experiments, except for Asn68, by alanine in the sequence used for modeling. This approach indeed resulted in a model with similar orientation of AtCYT6-NTD as compared to hCE and AtCYT6-CTD. However, AtCYT6-NTD was tilted relative to AtCYT6-CTD (Fig. 6A). Interestingly, in the model both the AtCYT6-NTD and -CTD presented a leucine residue in position P2‘, which perfectly matches the architecture of the S2’-binding site on AtLEGβ (Fig. S4B) (14).

**Figure 6:**
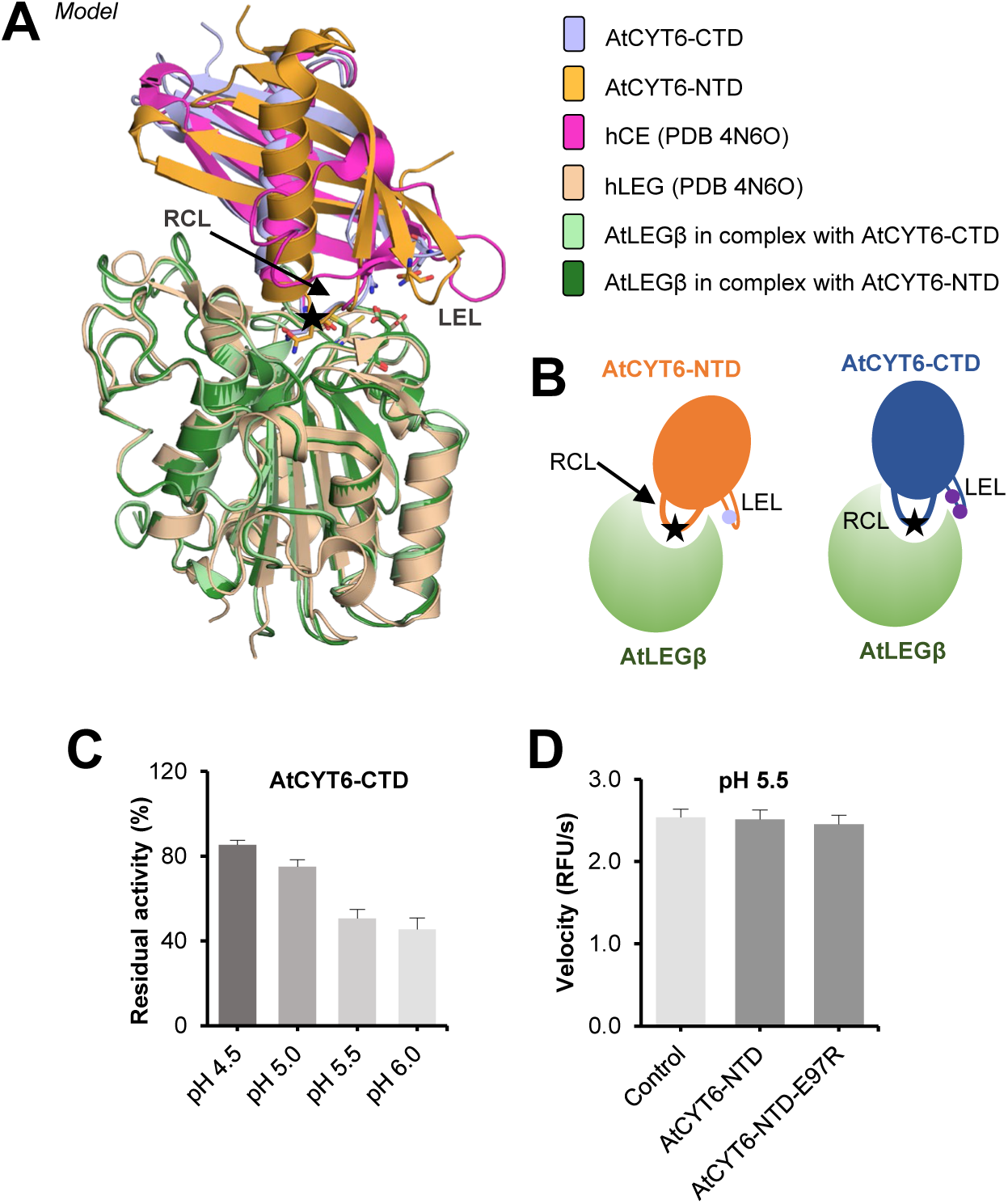
The interaction of AtCYT6 with AtLEGβ critically depends on the RCL and the LEL. **(A)** Proposed binding model of AtCYT6-NTD (orange) and AtCYT6-CTD (blue) to AtLEGβ (green) in comparison to the crystallographic structure (PDB 4N6O) of the complex between hLEG (wheat) and hCE (pink). The reactive center loops (RCLs) are indicated by a black arrow, the position of the P1-Asn residues are labeled with a star (LEL, legumain exosite loop). Models were generated with AlphaFold Multimer. **(B)** Model depicting the relative positions of Asn68/173 (black star) on the RCLs and Glu97 (light purple circle) and Arg205 and Lys208 (dark purple circles) on the LELs of AtCYT6-NTD or –CTD respectively. **(C)** Turnover of the AAN-AMC substrate by AtLEGβ at indicated pH values. The inhibition of AtLEGβ by AtCYT6-CTD is pH dependent. **(D)** AtCYT6-NTD and the AtCYT6-NTD-E97R mutant did not inhibit AtLEGβ at pH 5.5.

The legumain exosite loop (LEL) of hCE establishes an exosite interaction to hLEG, which turns hCE from a substrate into an inhibitor (21). Importantly, such a LEL structure was essentially missing in the AtCYT6 sequence. Instead, the LEL was replaced by a rather short hairpin loop in the AlphaFold model of AtCYT6 (Fig. 6A,C). Interestingly, the hairpin loop of AtCYT6-CTD harbored a double basic motif formed by Arg205 and Lys208 which, according to our model, could establish ionic interactions to Glu212 on AtLEGβ (Fig. 6A,B and S4,D). Glu212 is part of the S1‘-substrate binding site and sits directly next to the catalytic Cys211 residue. Based on this observation, we hypothesized that the Arg205-Lys208 double basic motif might serve as an electrostatic exosite anchor that stabilizes the enzyme-inhibitor complex, functionally resembling the LEL in hCE. To test this hypothesis, we prepared AtCYT6-CTD-R205A and -K208A mutants and tested their interaction with AtLEGβ. In line with our hypothesis, the mutants displayed reduced inhibition of AtLEGβ protease activity compared to the wild-type inhibitor (K_i AtCYT6-CTD-R205A_ = 3.36 ± 0.70 nM; K_i AtCYT6-CTD-K208A_ = 3.98 ± 0.66 nM) (Table 1). This experiment confirmed that the R205A-K208A double basic motif indeed functions as an important exosite that turns the AtCYT6-CTD into an inhibitor rather than a substrate. Consistent with the identified charged exosite interactions, we found that the inhibition of AtLEGβ by AtCYT6-CTD was pH-dependent. Inhibition was most efficient at around neutral pH and lower at very acidic pH (Fig. 6C). Moreover, we found that the processing of the different AtCYT6 constructs was also pH dependent. While the AtCYT6-CTD was rather resistant to cleavage at moderately acidic pH (pH 5.5), it was completely processed at pH 4.0, whereas the AtCYT6-NTD was quickly processed by AtLEGβ regardless of the pH (Fig. 5B,D). Importantly, the Arg205-Lys208 double basic motif was replaced by a negatively charged Glu97 on the AtCYT6-NTD (Fig. 6A,B, S4D). According to our model, Glu97 will result in repulsive interactions with Glu212 on AtLEGβ, which is in line with our observation that AtCYT6-NTD was not inhibiting AtLEGβ but rather acted as a substrate. To test whether we could turn AtCYT6-NTD into an inhibitor, we introduced an E97R mutation. Interestingly, we found that the AtCYT6-NTD-E97R mutant was not able to significantly impair AtLEGβ activity, even at a 10 : 1 inhibitor : enzyme ratio (Fig. 6D).

Sequence comparison of other five type II phytocystatins which had both domains individually tested against papain and different legumains indicated conservation of the properties seen in AtCYT6: (i) an asparagine residue on the RCL of N- and C-terminal domains, (ii) an R-G-X-K basic motif on the LEL of the legumain-inhibitory CTD, and a single Asp or Glu on the LEL of their NTDs (18,23,34) (Fig. S5).

Altogether, we conclude that complex formation between AtLEGβ and AtCYT6-CTD is pH-dependent, and its dissociation constant is modulated by the double basic motif Arg205-Lys208 within the LEL. Our experiments suggest that the LEL is not the only motif that is required for discrimination between inhibitory and non-inhibitory cystatin domains, as neither the R205A and K208A mutations on AtCYT6-CTD abolished legumain inhibition, nor the point mutation E97R elicited significant inhibition of AtCYT6-NTD against AtLEGβ.

### The AtCYT6-CTD has a pH-stabilizing effect on AtLEGβ

The catalytic domains of legumains are known to undergo irreversible denaturation at near neutral pH. Some interaction partners can stabilize the catalytic domain at such pHs, as is the case for hCE (21). Our model established a similar contact surface for AtLEGβ and AtCYT6-CTD to that of hCE and hLEG (Fig. 6A). Therefore, we hypothesized that a similar pH-protective role would be encoded in AtCYT6. To test this hypothesis, we performed nanoDSF experiments, which monitored the changes in intrinsic fluorescence of aromatic residues upon thermal unfolding. The ratio of fluorescence measured at 330 nm (absorption peak when aromatic residues are buried) and at 350 nm (absorption peak of solvent-exposed aromatic residues) was plotted against temperature, and the inflection temperatures (T_i_) of the samples were compared. As seen in Fig. 7A, at pH 6.5 AtLEGβ alone displayed a T_i_ of 49,6 °C, whereas when incubated with AtCYT6-FL the T_i_ shifted to 58.9 °C. Similarly, incubation of AtLEGβ with AtCYT6-CTD resulted in an increase of T_i_ to 57.0 °C. Taken together, complex formation resulted in thermal stabilization of AtLEGβ by approx. 8.5 °C. Importantly, when AtCYT6-NTD was incubated with AtLEGβ, no significant shift in T_i_ was detected (Fig. S6). Similarly, the P1 AtCYT6-CTD-N173A mutant was also not able to elicit the same stabilizing effect over AtLEGβ (Fig. 7B and S5B). However, the AtCYT6-CTD-K208A mutant showed a stabilization capacity comparable to its wild-type counterpart. This further confirmed that the stabilizing effect was directly linked to complex formation.

**Figure 7:**
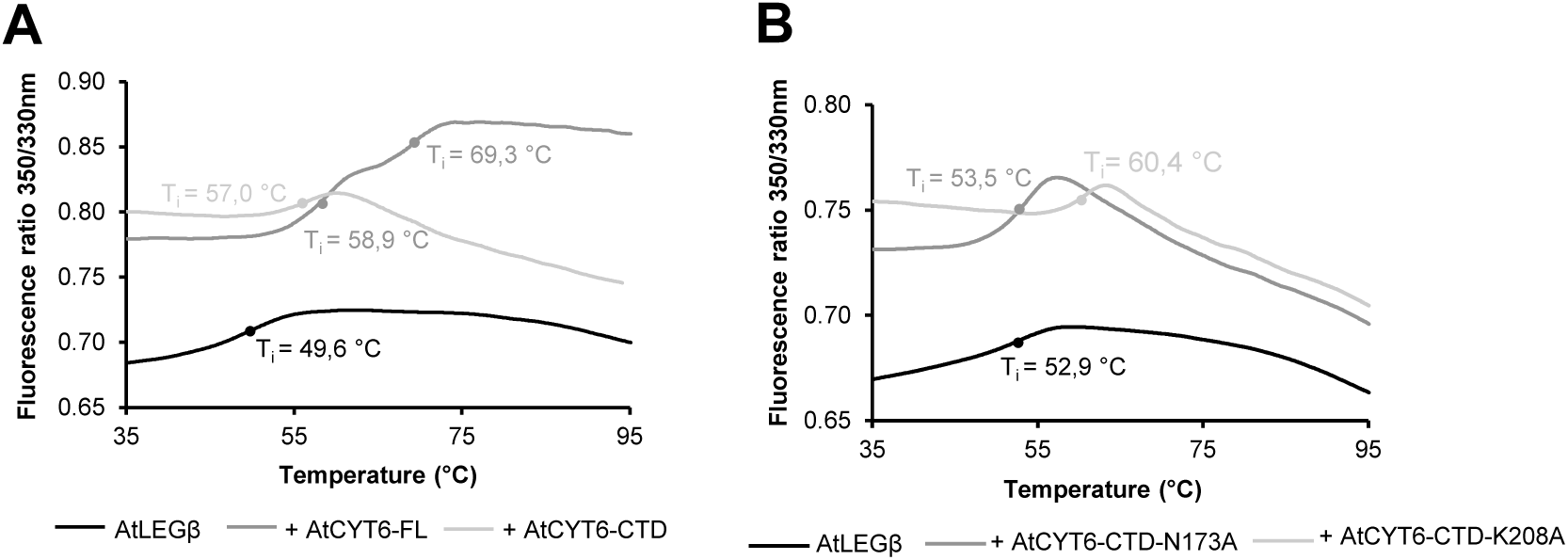
AtCYT6 has a stabilizing effect on AtLEGβ. The thermal stability of AtLEGβ was measured by nanoDSF (differential scanning fluorimetry) measurements. Unfolding transitions (inflection point; T_i_) are indicated by solid circles. **(A)** AtLEGβ was inhibited with MMTS and incubated at pH 6.5 in the absence (black) and presence of AtCYT6-FL (dark grey) or AtCYT6-CTD (light grey). **(B)** Same as **(A)** but after incubation with AtCYT6-CTD-N173A (dark grey) or AtCYT6-CTD-K208A (light grey). For control measurements of AtCYT6 constructs alone, see supplementary figure 6.

### Cystatin 6 is also a potent inhibitor of AtLEGγ

Once the structural requirements for inhibition of AtLEGβ by AtCYT6 were revealed, we wondered whether the inhibition would be specific to certain AtLEG isoforms. Considering that the catalytic domains of AtLEGβ and AtLEGγ are highly conserved and share critical features found to be important for inhibition by AtCYT6, e.g. the P1-Asn specificity and the Glu212 residue neighboring the catalytic cysteine, we hypothesized that AtLEGγ would also be inhibited by AtCYT6. Indeed, AtCYT6-FL and AtCYT6-CTD displayed a binding affinity (K_i_) of 1.21 ± 0.23 nM and 0.86 ± 0.22 nM, respectively, towards AtLEGγ (Table 2 and Fig. S7A,B), suggesting a physiological role for inhibition of both isoforms by AtCYT6. To test if the inhibition would be specific against plant legumains, we assessed the inhibition of human legumain by the different AtCYT6 constructs. Interestingly, we found significantly lower binding affinities for AtCYT6-FL and AtCYT6-CTD (K_i AtCYT6-FL_ = 106.4 ± 11.9 nM; K_i AtCYT6-CTD_ = 61.4 ± 7.5 nM), indicating intra-species specificity of AtCYT6 (Table 2 and Fig. S7D,E). The reduction in inhibition of human legumain might be explained by the missing S2’ substrate specificity pocket. Importantly, the N-terminal domain did not inhibit either of the enzymes (Fig. S7C,F).

**Table 2:** K_i_ values of different AtCYT6 constructs against AtLEGγ and human legumain (hLEG).

| | AtLEGγ ($K_i$ in nM) | hLEG ($K_i$ in nM) |
| --- | --- | --- |
| AtCYT6-FL | $1.21 \pm 0.23$ | $106.4 \pm 11.9$ |
| AtCYT6-NTD | n.i. | n.i. |
| AtCYT6-CTD | $0.86 \pm 0.22$ | $61.4 \pm 7.5$ |

## DISCUSSION

Based on our findings we suggest a bivalent binding mechanism of AtCYT6 to AtLEGβ which is established by two major binding sites. Binding site 1 mediates substrate-like binding via the P1-Asn173 residue on the RCL. Binding site 2 establishes electrostatic interactions via the double basic motif Arg205-Lys208 and thereby steers (pre-)complex formation (Fig. 8). Substrate-like binding inhibitors are not rarely substrates themselves – slow degradation may follow binding to the enzyme. Indeed, we observed that cleavage of AtCYT6-CTD upon incubation with AtLEGβ increased over time at pH 5.5 (Fig. 5B). We conclude that the inhibition of AtLEGβ by the AtCYT6-CTD happens at the expense of inhibitor degradation in a competitive, substrate-like manner. The presence of an asparagine residue in the corresponding RCL of non-inhibitory AtCYT6-NTD (Asn68) suggests that the P1-Asn is a prerequisite rather than a determining factor for AtLEG inhibition by phytocystatins.

**Figure 8:**
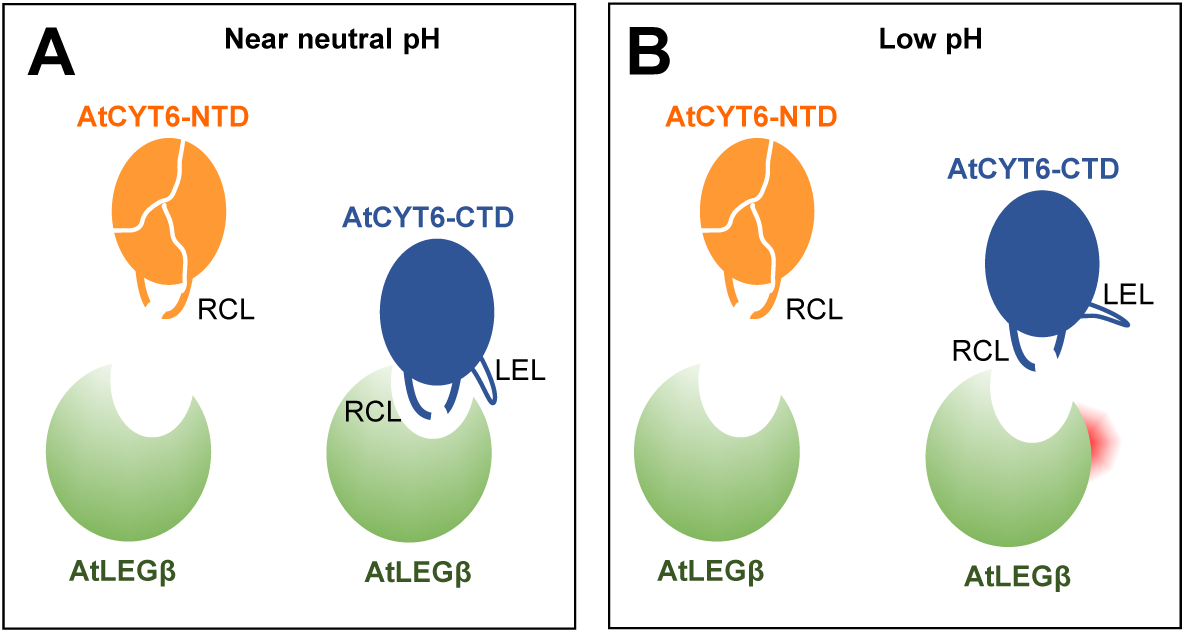
AtCYT6 is a substrate-like inhibitor of AtLEGβ. **(A)** At near neutral pH, the interaction of AtCYT6-CTD to AtLEGβ is stabilized by ionic interactions of the legumain exosite loop (LEL) to Glu202 on AtLEGβ. **(B)** At acidic pH (< 4.5) the complex is destabilized because of protonation of Glu202. Notably, AtCYT6-NTD functions as a substrate rather than an inhibitor at all pH conditions.

AtCYT6 is a high affinity inhibitor of legumains from *A. thaliana.* Despite the high conservation between the catalytic domains of AtLEGβ and AtLEGγ (more than 70% identity and 80% similarity), a 4-fold difference in K_i_ was observed for the AtCYT6-FL. This difference could be explained by a substitution on the non-prime side of the enzymes: a tyrosine in AtLEGβ (Tyr240) is substituted by a tryptophane (Trp249) in AtLEGγ, which results in a tighter S3 pocket to accommodate the bulky P3-Arg171. On the other hand, AtCYT6-FL features a 50-fold lower binding affinity to hLEG than it does for the *A. thaliana* legumains. In this case, the missing S2’ pocket in hLEG may be responsible for the reduction in affinity. Therefore, the P2’-Leu will bind with higher affinity to the *A. thaliana* legumains (35).

Our co-immunoprecipitation assays confirmed complex formation between AtCYT6-CTD and AtLEGβ *in vivo* (Fig. 4A,B). Notably, AtLEGβ was also recovered, although to a low amount, when anti-AtCYT6-NTD antibodies loaded with AtCYT6-NTD were used as a bait. This finding is in nice agreement with our observation that AtCYT6-NTD is recognized as a substrate by AtLEGβ rather than an inhibitor. Importantly, proteins like the PLCPs SAG12 and RD19 were exclusively co-eluted with anti-AtCYT6-NTD antibodies with and without recombinant AtCYT6-NTD. SAG12 is a senescence marker implicated in nitrogen content and yield of seeds of *A. thaliana*, whose expression is upregulated in *γvpe* knock-out plants (4,36,37).

We hypothesized that the LEL of the cystatins is a structural feature that discriminates inhibitory from non-inhibitory cystatin domains. However, introducing a supposedly ‘inhibitory’ AtCYT6-NTD-E97R mutation did not convert the non-inhibitory cystatin domain into a functional inhibitor. Our models also suggest that AtCYT6-NTD binds in a structurally different, tilted way to AtLEGβ. As a result, the AtCYT6-NTD LEL adopts a sterically different position, which can likely explain why the E97R mutation was not able to reconstitute inhibition on AtCYT6-NTD. Our models furthermore show that the orientations of Arg205 and Lys208 allow a strong electrostatic interaction with Glu212 from AtLEGβ – a neighbor residue to the catalytic cysteine Cys211. This double basic motif finds a resemblance in another human cystatin. Arg96 and Arg119 of the crystallographic structure of cystatin C (hCC, PDB 3GAX) align nicely with, and are oriented as Lys208 and Arg205 of our modeled AtCYT6-CTD (Fig. S8). Importantly, the interaction between hCC and hLEG is pH-sensitive; it is disrupted at pH 4.0. This exosite interaction is mediated via Glu190 on hLEG, which also sits directly next to the catalytic Cys189 (21). This is in good agreement with our experimental data and allows us to postulate that the LEL may be the structural motif that encodes pH-dependency to the stability of the enzyme-inhibitor complexes. However, since the R205A and K208A mutants showed a reduction in inhibition but not a complete loss of inhibition, we suggest that RCL cleavage and religation may be in an equilibrium at pH 5.5 and higher – but not at pH 4 – an important aspect for complex stabilization observed previously in hCE (38).

Human cystatin M/E is able to protect hLEG from quick and irreversible pH-driven denaturation. A pH-dependent stability was also observed for the catalytic domain of AtLEGβ: its conformational stability is high at pH 5.0 and decreases at pH 4.0 and 7.0 (14). Our results on the thermal unfolding (Fig. 7A) pose an interesting parallel, as the AtCYT6-CTD was able to significantly increase AtLEGβ stability at pH 6.5 and is, therefore, a candidate for legumain stabilization *in vivo*, e.g. outside of the vacuole.

Despite AtCYT6 being an AtLEGβ inhibitor, data from literature addressing their colocalization is still missing. The majority of AtCYT6 was shown to be expressed in seeds and siliques, and it is predicted to harbor an N-terminal signal peptide for extracellular secretion, as assigned in Uniprot (39,40). Our experiments revealed AtCYT6 to be an isoform-unspecific inhibitor displaying a high affinity for AtLEGβ and AtLEGγ. Along that line, CoIP experiments using immobilized anti-AtCYT6-CTD antibody incubated with *A. thaliana* leaf extract co-eluted AtLEGγ (data not shown). Interestingly, at pH 7.5 AtLEGγ was the sole legumain isoform that was co-eluted, whereas at pH 5.5 AtLEGβ was also present but in a significantly lesser extent. Together, these findings support the hypothesis that AtCYT6 could have tissue-specific targets. However, it remains unclear in which subcellular compartment and physiological context the inhibitor is to exert its inhibitory activity upon the enzymes. Despite the absence of experimental evidence of the presence of *A. thaliana* legumains in the extracellular environment, it is plausible to suppose its colocalization with its dedicated inhibitor, i.e., type II phytocystatins, in one or multiple physiological contexts. In that regard, the results of our pull-down assays in which AtLEGβ was used as a bait (Fig. 1D) show co-precipitation with a plasmodesmal protein (Uniprot O65449) and could indicate the presence of AtLEGβ in the extracellular environment (41). In this case, AtCYT6 could shield the enzyme from pH-driven denaturation upon alkalinization of the extracellular pH during plant-pathogen interaction and abiotic stress (42). Additionally, soluble extracellular content and vacuolar enzymes can meet up within the vacuolar system via endocytosis of apoplastic content (43–45). Lastly, the vacuole-mediated programmed cell death mechanisms comprise means by which AtCYT6 and AtLEGβ/γ may find each other. They consist of a series of events involving the lytic vacuole that culminate in either its collapse and consequent discharge of vacuolar proteases into the or fusion with the cell membrane, which also promotes mixing of vacuolar and apoplastic content (6,46–48).

Our experimental data demonstrate AtCYT6-NTD to be a potent inhibitor of papain, and AtLEGβ to be capable to promptly process it, and its equal competence to process AtCYT6-CTD at acidic pH (Fig. 5B,D). It is tempting to speculate that such post-translational processing of AtCYT6 could release the individual N- and C-terminal domains as truncation products, which would consequently allow them to be translocated into different subcellular compartments *in vivo*. This hypothesis is further strengthened by studies on type III phycys. Upon incubation with multiple endoproteases, functional subdomains with sizes of approximately 10, 22 and 32 kDa were released from the 85 kDa potato multicystatin, and retained their inhibitory activity against papain (49). Similarly, tomato leaves expressed a multicystatin of approximately 88 kDa which released functional fragments after short-term proteolytic processing by trypsin and chymotrypsin (50).

Taken together, our work suggests that the separation of functional subdomains within multi-domain phytocystatins could be a feature conserved throughout the cystatins. Additionally, the processing of AtCYT6 *in vitro* suggests a regulatory mechanism of its inhibitory activity by itself, as it demonstrates that individual domains can be processed by selected enzymes under differential conditions such as pH.

## Material & Methods

### Construct design

Expression constructs of full-length *Arabidopsis thaliana* cystatin 6 (AtCYT6-FL) were designed based on the sequence deposited in Uniprot under the accession code Q8HOX6-2. A full-length cDNA sequence lacking the N-terminal signal peptide (M1-M34) and harboring 5’ NdeI and 3’ XhoI restriction enzyme cleavage sites was purchased from Eurofins Genomics (Ebersberg, Germany). The expression construct was subcloned into the pET-22b(+) expression vector using the same restriction enzymes, resulting in a sequence encoding a C-terminal His_6_-tag. Additionally, the expression construct was also subcloned into the pET-15b vector, to obtain an N-terminal His_6_-tag. For that, a PCR reaction was performed using the primers TATAGGTACCGCTTTAGTCGGAGGTGTTGGCG (forward, KpnI recognition site) and GGTGGGATCCTCAGTCATGGTGTTGCTCCGCGTGG (reverse, BamHI). The PCR product was purified, digested by the respective restriction enzymes, and ligated into the pET-15b vector. Furthermore, constructs composed of the individual C-terminal domain (AtCYT6-CTD) and the C-terminal domain harboring the interdomain linker (AtCYT6-CTD_long_) were prepared using a similar protocol. For the AtCYT6-CTD CAAGGGTACCGAACATGAATCTGGATGGAGGG (forward, KpnI) and the full-length reverse primer were used, and for the AtCYT6-CTD_long_, a TGCCGGTACC GCCCCTGCTATCACTTCC TCCG (forward, KpnI) forward primer was used together with the reverse primer of the full-length construct. As the expression of the AtCYT6-NTD construct in the pET15b vector was inviable, we cloned the sequence into vector pET-28 and it included, as a result, a C-terminal His_6_-tag. A PCR reaction with the forward primer AGTCCCATGGCTTTAGTCGGAGGTGTTGGCG (NcoI) and reverse primer AGCTCTCGAGATCGGAGGAAGTGATAGCAGGGGC (XhoI) was done, and followed by purification, enzymatic digestion, and ligation into pET28. AtCYT6 point mutants were prepared using round-the-horn site directed mutagenesis. The forward primer GCGTCCTTGTTCCCTTATGAACTTCTGGAGGTTGTGC and reverse primer AGACCTCTGCTGAATGGTCTTGACAGCCTGCTCAGC were used to obtain the P1-Asn mutant N173A. The LEL mutation R205A in AtCYT6-CTD was obtained with the forward GCGGGAGAAAAAGAGGAAAAGTTCAAGGTGGAAGTTCAC and reverse CTTCAACTTCAGTAGCATGTTGTATTTTGCAGCCTCGC primers. The K208A mutation was introduced in AtCYT6-CTD with the forward GCGGAGGAAAAGTTCAAGGTGGAAGTTCACAAGAACC and reverse TTCTCCCCTCTTCAACTTCAGTAGCATGTTGTATTTTGC primers, and E97R was introduced in AtCYT6-NTD with CGCGCAGGACAGAAGAAGCTATACGAAGC (forward) and CAGAATCTCCAGAGTCAGGTGATGC (reverse) primers.

### Expression

The final expression vectors were transformed into the *E. coli* BL21(DE3) strain for expression. Expression flasks of 2 L containing 500 mL of LB medium (Carl Roth LB-Medium, Lennox) and ampicillin (for constructs cloned into pET15b) or kanamycin (for constructs cloned into pET28) at 100 μg/ml concentration were inoculated with 25 ml of preculture and incubated at 37 °C and 240 rpm until an OD_600_ of approximately 1.0 was reached. Expression was induced upon addition of IPTG (Isopropyl β-D-1-thiogalactopyranoside) at a final concentration of 1 mM. After induction, the cultures were kept overnight at 25 °C and 240 rpm. The cells were harvested and either lysed for immediate use, or frozen as pellets at -20 °C.

### Purification

Cell pellets were resuspended in 25 ml of lysis buffer (Tris-HCl 100 mM pH 7.5, NaCl 500 mM) and lysed by sonication at 40 % power, 4 times for 45 seconds (Bandelin Sonopuls) in the presence of EDTA-free, EASY-pack protease inhibitor cocktail tablets (Roche, Basel) according to instructions of the manufacturer. The lysates were centrifuged at 17500 g for 1 h at 4 °C. The supernatant was incubated for 30 min at 4 °C with 5 mL of Ni^2+^-beads (Qiagen Ni-NTA Superflow) equilibrated in lysis buffer. After removal of the flow-through, the beads were washed with 2 column volumes of buffer (Tris-HCl 100 mM pH 7.5, NaCl 500 mM) containing 5 mM, 10 mM and 20 mM imidazole, in that order. The elution of the protein of interest was done in a buffer composed of 100 mM Tris-HCl pH 7.5, 500 mM NaCl, and 250 mM imidazole in 3 steps of 1 column volume each with 15 min incubation. The eluate was concentrated in Amicon Ultra centrifugal filter units (MWCO: 3 kDa; Cytiva). Following affinity chromatography, the proteins were further purified by size exclusion chromatography using an ÄKTA FPLC system equipped with an S200 10/300 GL or S75 10/300 GL column equilibrated in a buffer composed of 20 mM Tris-HCl pH 7.5, 50 mM NaCl and 2 mM DTT (if indicated).

### Preparation of *Arabidopsis thaliana* legumain-β (AtLEGβ) and -γ (AtLEGγ), human legumain and papain

Papain from *Carica papaya* was purchased from Merck (Darmstadt, Germany). AtLEGβ, AtLEGβ-C211A, AtLEGγ and human legumain were expressed, purified and activated based on a protocol previously described (51). Briefly, the sequence of corresponding constructs of proLEG were cloned into pLEXSY-sat2.1 vectors and transfected into LEXSY P10 host strain of *Leishmania tarentolae* cells of the LEXSYcon2.1 expression system (Jena Biosciences, Jena, Germany). The resulting constructs carried N-terminal His_6_-tags and an N-terminal signal sequence for secretion into the supernatant. Cultures were grown in brain heart infusion (BHI) medium (Jena Biosciences) in tissue culture flasks with 5 μg/ml of hemin, 50 units/mL penicillin and 50 units/mL streptomycin (Pen-Strep, Carl Roth) in the presence of nourseothricin (Jena Bioscience) as selection antibiotic. Expression was carried out for 48 hours in the dark at 26 °C and 140 rpm. The supernatant containing the proteins of interest were harvested by centrifugation and incubated with Ni^2+^-beads at 4 °C. The washing buffer consisted of 50 mM HEPES pH 7.5 and 300 mM NaCl. The elution buffer was the same as washing buffer with addition of 250 mM imidazole. After elution, the proteins were concentrated, and PD-10 columns (GE Healthcare, Uppsala, Sweden) were used for buffer exchange with storage buffer 20 mM HEPES pH 7.5 and 50 mM NaCl. At this point, proteins were either stored at -20 °C for further use or auto activated for elimination of the C-terminal prodomain. The pH-driven auto activation was performed with incubation at room temperature at pH 3.5 - 4.0 after which size exclusion chromatography or buffer exchange with PD-10 columns was performed with buffer composed of 20 mM citric acid pH 4.0 - 5.0 and 50 mM NaCl.

The details described above may vary slightly according to the legumain isoform in question. For a more detailed and precise protocol, please refer to the respective publications (13,14,52).

### Differential scanning fluorimetry

10 μL of AtLEGβ (1 mg/ml) in a buffer composed of 20 mM citric acid buffer pH 4.0, 50 mM NaCl, and 0.5 mM MMTS – methyl methanethiosulphonate) were mixed with an equimolar amount of cystatins for 5 min at 21 °C, in a final volume of 12 μL. Control reactions were supplemented with buffer instead of cystatins or legumain, respectively. For measurements done at pH 6.5, 1 μL of 1 M MES pH 6.5 buffer was added immediately before the addition of the cystatin. Thermal denaturation curves were obtained using the Nanotemper Tycho NT.6 instrument by monitoring the fluorescence intensity at 330 and 350 nm upon heating the samples form 35 °C to 95 °C.

### Activity assays

Initial velocities were obtained by monitoring the fluorescence increase (in RFU) following the breakdown of synthetic peptidic substrates in an Infinite M200 Plate Reader (Tecan) at 25 °C. For legumain assays, the Z-Ala-Ala-Asn-AMC (Bachem) substrate was used at 50 µM concentration in reaction buffer composed of 20 mM citric acid pH 5.5, 100 mM NaCl, 0.02 % Tween 20, and 2 mM DTT). For papain assays, the Z-Phe-Arg-AMC (Bachem) substrate was at 50 µM concentration in reaction buffer composed of 20 mM MES pH 6.0, 50 mM NaCl, 0.02 % Tween 20, and 2 mM DTT). The enzymes and the inhibitors were added 10-fold concentrated in relation to their desired final concentration in the assay. Control experiments were supplemented with buffer instead of inhibitor. Reactions were started by addition of the enzyme. The first 100 seconds of the reactions were used for the calculation of the initial velocities. For the K_i_ determinations, the final concentration of enzymes was 50 nM, and the inhibitor concentrations were defined by 1:2 serial dilutions starting at 15-fold the enzyme concentration, measured in quadruplicates. To obtain Ki values, data points were fitted to the Morrison equation using the GraphPad Prism program (version 5.0, La Jolla, CA).

### Analysis of complex formation via co-migration assays

Co-migration experiments were performed in buffer composed of 20 mM MES pH 6.0, and 100 mM NaCl. 400 μg of AtLEGβ or papain were inhibited with 0.5 mM or 5 mM S-methyl methanethiosulfonate (MMTS, Merck), respectively. Subsequently, an equimolar amount of AtCYT6-derived constructs was added, and enzyme and inhibitor were incubated for 1 h on ice. Samples were loaded on a Superdex S75 10/300 GL column (Cytiva) preequilibrated in assay buffer at 21 °C. Immediately after elution, MMTS was added at a final concentration of 5 mM. Samples collected were analyzed by SDS-PAGE. Samples containing legumain were subjected with 1 mM of the covalent Ac-Tyr-Val-Ala-Asp-chloromethyle ketone (YVAD-cmk) inhibitor prior to the addition of the loading buffer.

### pH-dependent cleavage

AtLEGβ was incubated with AtCYT6-derived constructs at a 1:5 molar ratio in a buffer composed of 100 mM citric acid pH 4.0 or 5.5, 100 mM NaCl, and 2 mM DTT at 21 °C. Samples were collected after 1, 5 and 10 minutes of incubation and transferred to a new tube containing 0.1 μL of YVAD-cmk inhibitor (final concentration: 1mM). Subsequently, samples were analyzed by SDS-PAGE.

### Cleavage analysis by mass spectrometry

AtCYT6-derived constructs were incubated with AtLEGβ in a 1:5 molar ratio in a buffer composed of 100 mM citric acid pH 4.0 or 5.5, 100 mM NaCl, and 2 mM DTT at 21 °C for 30 minutes. For mass spectrometric analysis, samples were desalted with C_18_ ZipTips (Merck Millipore), eluted from the tips with 50 % acetonitrile in 0.1 % formic acid and directly infused into the mass spectrometer (Q-Exactive, Thermo Fisher Scientific) at a flow rate of 1 µl/min. Capillary voltage at the nanospray head was 2 kV. Raw data were processed with Protein Deconvolution 2.0 (Thermo Fisher Scientific). Masses were assigned to the protein sequence with the Protein/Peptide Editor module of BioLynx (part of MassLynx V4.1, Waters).

### Modelling of protein complexes

The models of the complexes were obtained using Alphafold Multimer (via Colab notebook). The sequence used for modeling of AtLEGβ comprised residues V48-N329 according to Uniprot accession code Q39044. The models of AtCYT6-NTD and AtCYT6-CTD corresponded to residues A35-D127 and G141-D234, respectively, from the Uniprot code Q8H0X6. The PyMOL Molecular Graphics System (Schrödinger, LLC) was used to illustrate protein models.

### Pull-down assay with recombinant AtLEGβ

For pulldown experiments with recombinant AtLEGβ, VPE0 mutant (accession N67918) seed stocks were obtained from the Nottingham Arabidopsis Stock Center (NASC, Nottingham, UK) and grown on soil at short day conditions (9 h light with an intensity of 120 μE m−2 s−1 at 22 °C and 15 h darkness at 18 °C, 75 % RH) after stratification for 3d at 4 °C. Leaves of 5-week-old plants were harvested and homogenized using a Polytron PT-2500 (Kinematica, Luzern, Switzerland) in extraction buffer (100 mM MES pH 6.0, 100 mM NaCl, 250 mM sucrose, 1 % Triton X-100, and HALT protease inhibitor cocktail; ThermoFisher, Dreieich, Germany) followed by centrifugation at 4000 g, 4 °C for 5 minutes. The lysate was then quantified using the BCA assay (ThermoFisher, Dreieich, Germany). A total of 2 mg of lysate per condition together with 25 μL of Dynabeads^TM^ for His-Tag isolation (Invitrogen) were incubated with His_6_-AtLEGβ or the His_6_-proAtLEGβ-C211A dead mutant or lysate alone for an hour at room temperature with rotation. To block its proteolytic activity, His_6_-AtLEGβ was pre-inhibited with MMTS. The Dynabeads were then washed four times with 400 μL of wash buffer (100 mM MES pH 6.0, 300 mM NaCl, 1 % Triton X-100). Elution was performed in 100 μL of elution buffer (300 mM imidazole, 300 mM NaCl, 100 mM HEPES pH 7.5, 1 % Triton X-100).

For sample processing prior to mass spectrometry measurement, each eluted fraction was then heated to 56 °C for 10 minutes followed by reduction with 20 mM DTT for a further 30 minutes under shaking conditions at 1500 rpm and alkylation with 50 mM CAA (chloroacetamide) at room temperature for 30 minutes in the dark. The reaction was quenched with 50 mM DTT for 30 minutes and purified with SP3 beads. For each reaction, 5 μL of SP3 bead stock (1:1 mixture of hydrophilic and hydrophobic carboxylated beads, 1 µL per 20 µg of protein) was added and protein binding to the beads was induced with a final concentration of 80 % ethanol followed by incubation at room temperature for 30 minutes with rotation. The beads were then washed twice with 90 % acetonitrile before being resuspended in trypsin solution (50 mM HEPES pH 7.5, 5 mM CaCl_2_, 1 μg trypsin) and incubated at 37 °C for 16 hours at 1500 rpm. For triplex differential labelling of the peptides, the conditions were labelled with 30 mM light, CH_2_O (lysate alone), medium, CD_2_O (AtLEGβ), heavy, ^13^CD_2_O (proAtLEGβ-C211A) and 15 mM sodium cyanoborohydride at 37 °C for one hour, 1500 rpm. The reaction was then quenched with 100 mM Tris for 30 minutes. The labelled peptides from the three conditions were pooled 1:1:1 and acidified with final 1 % formic acid trifluoroacetic acid before passing through pre-activated C18 StageTips.

### AtCYT6 CoIPs

For co-immunoprecipitation assays we purchased rabbit sera (BioGenes, Berlin, Germany) of animals treated with injections of AtCYT6-NTD or AtCYT6-CTD separately. The antibodies raised against each AtCYT6 domain were purified from the sera using CNBr-activated Sepharose 4 Fast Flow beads (GE Healthcare, Uppsala, Sweden) with immobilized recombinant AtCYT6-NTD and AtCYT6-CTD according to the manufacturer’s protocol. The integrity of the purified antibodies was tested in western blot assays against AtCYT6-NTD and AtCYT6-CTD, and each pool of purified polyclonal antibodies displayed high sensitivity and specificity to its respective antigen after this step, so we moved on to the co-immunoprecipitation assay with plant material. To that end, we used rec-Protein G-Sepharose 4B Conjugate beads (Invitrogen, Camarillo, USA) cross-linked to the polyclonal antibodies with dimethylpimelidate-dihydrochlorid (Thermo Fisher Scientific) to prevent leakage of antibodies to the supernatant in further steps. The empty beads, i.e. Sepharose beads conjugated with protein G that were not cross-linked to antibodies, constituted an important control and are referred as *bait 1*. The beads that were cross-linked to antibodies are *bait 2*. For the co-immunoprecipitation using antibodies loaded with their recombinant antigen as baits, antibodies freshly cross-linked to the beads were further incubated with recombinantly expressed antigens, i.e. AtCYT6-NTD or AtCYT6-CTD, in buffer composed of 50 mM HEPES pH 7.5 and 100 mM NaCl for 1 h at 4 °C. The unbound antigens were washed away with multiple centrifugations (at 4,000 g at 4 °C) with elimination of the supernatant and resuspension in buffer HEPES 50 mM and NaCl 100 mM pH 7.5 until the UV signal at 280 nm was stable and null. This corresponds to the *bait 3*.

The preparation of the plant material followed with the maceration of 100 mg of *A. thaliana* seeds in liquid nitrogen until a fine powder was obtained. Immediately after grinding, 0.6 mL of incubation buffer (citric acid 100 mM and NaCl 250 mM pH 5.5 with inhibitor cocktail (Roche, Rotkreuz, Switzerland) dilution according to the instructions of the manufacturer)) was added and thoroughly mixed for 10 minutes at 4 °C. The suspension was centrifuged three times at 13,000 g for 15 minutes at 4 °C with change of tubes in between. The resulting supernatant was split in three tubes with the same volume of material, and to each tube either bait 1, 2 or 3 was added and incubated at 4 °C with gentle and constant rotation for 1 hour. After that, the beads were washed thoroughly with incubation buffer in multiple cycles of centrifugation and discard of supernatant until the UV signal at 280 nm of the supernatant was stable and null. The co-precipitated proteins were eluted with 1x Laemmli/SDS sample buffer without DTT at 37 °C for 15 min in a shaker with 750 rpm. Then, we performed an in-gel digestion with trypsin. The eluted samples from co-immunoprecipitation assays with bait 1, 2 and 3 were loaded onto an acrylamide gel for SDS-PAGE and allowed to run 1 - 2 cm into the separation gel. All the area supposedly containing proteins was cut out, reduced to pieces of approximately 1 mm x 1 mm and collected in low binding tubes. The gel pieces of each tube were washed with buffer A (50 mM ammonium bicarbonate) and then with buffer B (25 mM ammonium bicarbonate in 50 % acetonitrile). The proteins trapped within the gel pieces were reduced with reduction buffer (buffer A with 20 mM DTT) for 30 min at 56 °C, alkylated with alkylation buffer (buffer A with 10 mM iodoacetamide) for 30 min at room temperature in the dark, and then washed with buffer A and B. The drying of the gel pieces started with shrinking by addition of 100% acetonitrile and finished with complete drying in a SpeedVac at 50 °C until the gel pieces were fully dried. For trypsin digestion, the gel pieces were hydrated again with a solution of 12.5 ng/μL trypsin in digestion buffer (25 mM ammonium bicarbonate, 5 % acetonitrile and 5 mM CaCl_2_) and incubated overnight at 37 °C. The supernatants containing the peptides were transferred to new tubes and 100 % acetonitrile was added to the gel pieces and incubated for 15 minutes at room temperature. The acetonitrile supernatant was combined with the supernatant previously collected and the resulting sample was dried in a SpeedVac at 50 °C until a minimum volume was reached. The volume of the samples was adjusted to 25-50 μL and pH adjusted to < 3 with 2.5 % formic acid before desalting the samples with C18 double-layer stage tips (Empore Octadecyl C18, Supelco, USA).

### Western blots of CoIP samples

The samples collected in the CoIP experiment were further analyzed via Western blotting using rabbit anti-AtCYT6-NTD or -CTD antisera respectively. Specifically, 2 µl of the elutions were subjected to SDS-PAGE and transferred to an Amersham Protran 0.45 μm nitrocellulose blotting membrane (GE Healthcare, Uppsala, Sweden) using a Trans-Blot semi-dry transfer cell (Bio-Rad, Vienna, Austria). 10 ng of recombinant AtCYT6-FL, -NTD and -CTD were loaded as controls. The membranes were washed 3 times with TBST and subsequently blocked with 5 % milk powder dissolved in TBST. Incubation with the respective rabbit antisera was done in the dark at 22 °C for 1 hour in a 1:40 dilution with TBST. Subsequently, membranes were washed with TBST and incubated with goat anti-rabbit-IgG coupled to horseradish peroxidase (Santa Cruz Biotechnology, Dallas, USA) in a 1:12000 dilution in TBST for 1 hour. Chemiluminescence signals were detected using the Image Lab software (version 6.0.1; Bio-Rad, Vienna, Austria) after addition of Amersham ECL Prime Western Blotting detection reagent (GE Healthcare, Uppsala, Sweden).

### Mass spectrometry analysis of pulldown and CoIP samples

AtLEGβ pulldown assays were analyzed on a two-column nano-HPLC setup (Ultimate 3000 nano-RSLC, Thermo) equipped with Acclaim PepMap 100 C18 columns (ID 75 μm, particle size 3 μm columns, trap column of 2 cm length, analytical column of 50 cm length, Thermo) operated at 60°C. For AtCYT6-coIP samples, the same system was equipped with a µPAC reverse phase trap and analytical (50 cm) columns (Thermo) operated at a column temperature of 40 °C. Peptides were eluted with a binary gradient from 5–32.5% B for 80 min (A: H2O + 0.1% FA, B: ACN + 0.1% FA) and a total runtime of 2 h per sample. Spectra were acquired in data-dependent data acquisition mode using a high-resolution Q-TOF mass spectrometer (Impact II, Bruker, Bremen, Germany) as described (53).

### Mass spectrometry data analysis

Tandem mass spectra acquired for AtLEGβ pulldown assays were analyzed with the MaxQuant software package (54), v1.6.0.16, with the Uniprot *Arabidopsis thaliana* reference proteome database (release 2019_03) with appended sequence of recombinant proAtLEGβ-C211A and list of common contaminants included in MaxQuant. Searches considered tryptic digest with up to 2 missed cleavages, Met oxidation and protein N-terminal acetylation as variable modifications and triplex stable isotope labeling of peptide N-termini and Lys residues with light formaldehyde and sodium cyanoborohydride (+28.0313 Da), deuterated formaldehyde (^12^CD_2_O) and sodium cyanoborohydride (+32.0564 Da) and ^13^CD_2_O formaldehyde and sodium cyanoborodeuteride (+36.0756 Da). Standard settings for Bruker QTOF data were used with standard settings for protein identification including PSM and protein FDR<0.01. AtCYT6-CoIP data was analysed with MaxQuant v.2.0.3.0 using essentially the same settings and database, except that no labeling was applied and LFQ was enabled with minimal LFQ count was set to 1 for label-free quantification.

### Alignments

All the alignments were done with Clustal Omega (55). The alignment depicted in Fig. 1 comprises the sequences of hCE/M, Uniprot entry Q15828 lacking N-terminal signal peptide, and AtCYT6, Uniprot entry Q8H0X6 lacking the N-terminal signal peptide, and was further illustrated with the protein sequence editor Aline (56). The other alignments were done with the full-length sequences of cystatins 1-7 of *A. thaliana* (AtCYT1 (Q945Q1), AtCYT2 (Q8L5T9), AtCYT3 (Q41906), AtCYT4 (Q84WT8), AtCYT5 (Q41916), AtCYT6 (Q8H0X6), AtCYT7 (Q8LC76)), cystatin 12 from *Oryza sativa*, (OsCYT12, Q0JNR2), cystatin 4 from *Hordeum vulgare* (HvCYT4, Q1ENE9), cystatin 1 from *Fragaria ananassa* (FaCYT1, Q4GZT8) and cystatin 1 from *Fragaria chiloensis* (FcCYT1, A0A2U9AEI9).

### Data availability

The mass spectrometry proteomics data have been deposited to the ProteomeXchange Consortium (57) via the PRIDE (58) partner repository with the dataset identifiers PXD042260 for the AtLEGb pulldowns (username; password: ZGuPv4w7) and PXD042261 for the AtCYS6 coIP data (username; password: lDPtFrnJ).

## Acknowledgements

The authors wish to thank Martina Wiesbauer for technical assistance and Melissa Mantz for mass spectrometry co-IP data acquisition.

## Funding and additional information

This work was supported by the Austrian Science Fund (FWF, project number P31867, to E.D.).

