## Supporting Information for "Phytocystatin 6 is a context-dependent, tight-binding inhibitor of *Arabidopsis thaliana* legumain isoform β"

**Affiliations:**

### Supplementary Figures

```
AtCYT-2     -------MATM--LKVSLVLSLLGFLVIAVVTPSAANPFRKSVVLGGKSGVPNIRTN-RE     50
AtCYT-7     MDMRRASMCMMLICVSLVLLSGFGQFVICSEEKGTYNDNVVKMKLGGFSDSKNDWNGGKE     60
AtCYT-1     --------------------------------MAD---QQAGTIVGGVRDIDANAND-LQ     24
AtCYT-3     --MESKT-----FWIVTLLLCGTIQLAICRSEEKS---TEKTMMIGGVHDLRGNQNS-GE     49
AtCYT-6     -MMRSRFLLFIVFFSLSLFISS----LIASDLGFC---NEEMALVGGVGDVPANQNS-GE     51
AtCYT-4     ------MMMKSLICLSLILLPLVSVV----------EGLGGGGGIGSRKPIKNV--SDPD     42
AtCYT-5     --MTSKVVFLLLLSLVVVLLPLYA---------------SAAARVGGWSPISNV--TDPQ     41
                                                        :*.           :

AtCYT-2     IQQLGRYCVEQFNQQAQNEQGNIGSIAKTDTAISNPLQFSRVVSAQKQVVAGLKYYLRIE     110
AtCYT-7     IDDIALFAVQEHNRRE-----------------NAVLELARVLKATEQVVAGKLYRLTLE     103
AtCYT-1     VESLARFAVDEHNKNE-----------------NLTLEYKRLLGAKTQVVAGTMHHLTVE     67
AtCYT-3     IESLARFAIQEHNKQQ-----------------NKILEFKKIVKAREQVVAGTMYHLTLE     92
AtCYT-6     VESLARFAVDEHNKKE-----------------NALLEFARVVKAKEQVVAGTLHHLTLE     94
AtCYT-4     VVAVAKYAIEEHNKES-----------------KEKLVFVKVVEGTTQVVSGTKYDLKIA     85
AtCYT-5     VVEIGEFAVSEYNKRS-----------------ESGLKFETVVSGETQVVSGTNYRLKVA     84
            :  :. :.:.:.*:.                   .  *    :: .  ***:*  : * :

AtCYT-2     VTQPNGSTRMFDSVVVIQPWLHSKQLLGFTPVVSPVY-----------------------     147
AtCYT-7     VIEAG-EKKIYEAKVWVKPWMNFKQLQEFKNI--IPSFTISDLGFKPDGNGFDWRSVSTN     160
AtCYT-1     VADGE-TNKVYEAKVLEKAWENLKQLESFNHLHDV-------------------------     101
AtCYT-3     AKEGD-QTKNFEAKVWVKPWMNFKQLQEFKESSS--------------------------     125
AtCYT-6     ILEAG-QKKLYEAKVWVKPWLNFKELQEFKPASDAPAITSSDLGCKQGEHESGWREVPGD     153
AtCYT-4     AKDGGGKIKNYEAVVVEKLWLHSKSLESFKAL----------------------------     117
AtCYT-5     ANDGDGVSKNYLAIVWDKPWMKFRNLTSFEPANNGRFL----------------------     122
              :     : : :  *  : * : :.*  *

AtCYT-2     ------------------------------------------------------------     147
AtCYT-7     NPEVQEAAKHAMKSLQQKSNSLFPYKLIDIILARAKVVEERVKFELLLKLERGNKLEKFM     220
AtCYT-1     ------------------------------------------------------------     101
AtCYT-3     ------------------------------------------------------------     125
AtCYT-6     DPEVKHVAEQAVKTIQQRSNSLFPYELLEVVHAKAEVTGEAAKYNMLLKLKRGEKEEKFK     213
AtCYT-4     ------------------------------------------------------------     117
AtCYT-5     ------------------------------------------------------------     122

AtCYT-2     ---------------------     147
AtCYT-7     VEVMKDQTGKYE---------     232
AtCYT-1     ---------------------     101
AtCYT-3     ---------------------     125
AtCYT-6     VEVHKNHEGALHLNHAEQHHD     234
AtCYT-4     ---------------------     117
AtCYT-5     ---------------------     122
```

**Supplementary figure 1: Sequence alignment of the seven predicted phycys isoforms of *A. thaliana*.** Sequences were retrieved from Uniprot entries Q945Q1 (AtCYT1), Q8L5T9 (AtCYT2), Q41906 (AtCYT3), Q84WT8 (AtCYT4), Q41916 (AtCYT5), Q8H0X6 (AtCYT6) and Q8LC76 (AtCYT7). Motifs for inhibitory activity against papain and legumain are highlighted in solid black boxes and dashed black box, respectively.

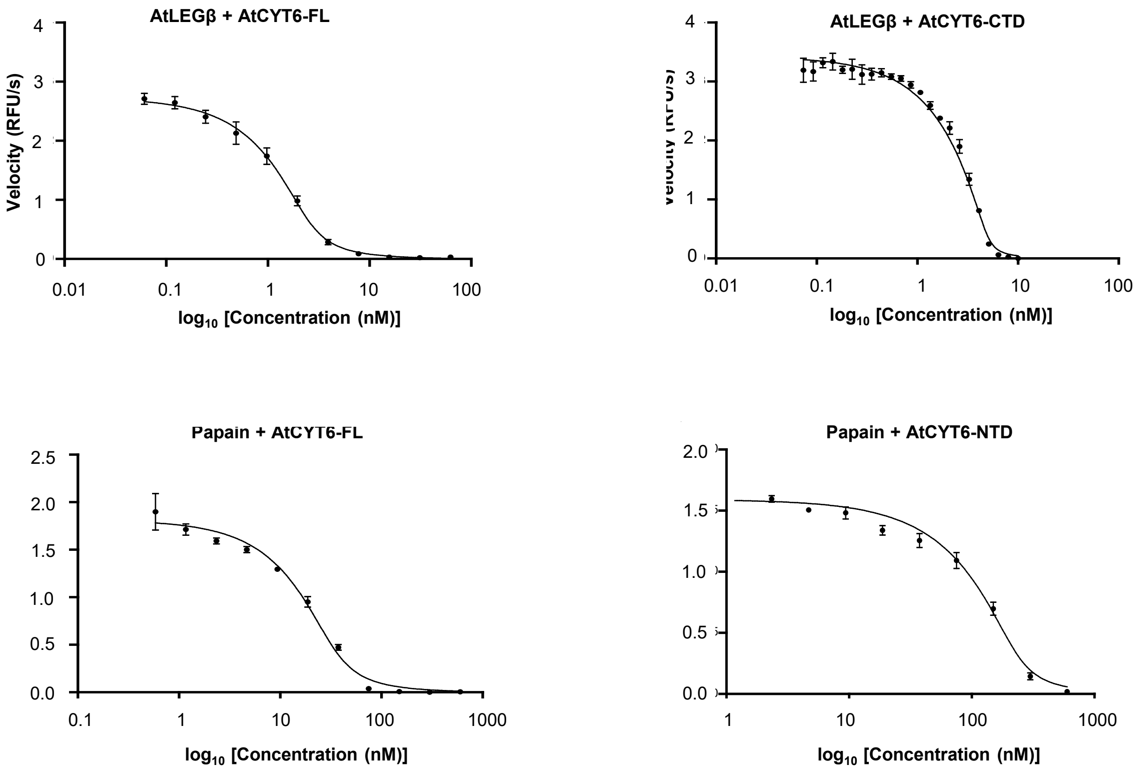


**Supplementary figure 2: The *K*i of indicated AtCYT6 constructs towards AtLEGβ and papain was determined using Morrison's equation.** **(A)** and **(B)** Activity was measured at pH 5.5 using the AAN-AMC substrate. **(C)** and **(D)** Activity was measured at pH 6.5 using the FR-AMC substrate.

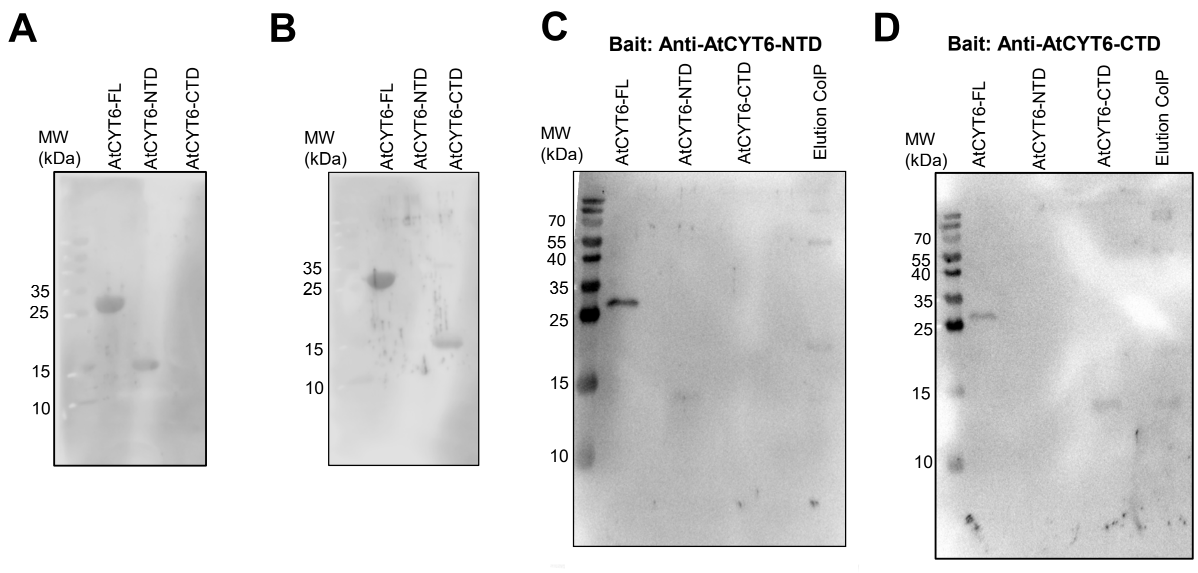


**Supplementary figure 3: Western-blot experiments showed that AtCYT6 was processed in *A. thaliana* seed extract. (A)** Western blot against indicated recombinant AtCYT6 proteins using anti-AtCYT6-NTD antiserum as primary antibody. **(B)** Same as **(A)** but using anti-AtCYT6-CTD antiserum. **(C)** A seed extract was prepared at pH 5.5 and co-immunoprecipitated with immobilized anti-AtCYT6-NTD antibody as a bait. The elution fraction was analyzed by western blotting using anti-AtCYT6-NTD antibodies as primary antibody. **(D)** Same as **(C)** but using anti-AtCYT6-CTD as bait in the CoIP and anti-AtCYT6-CTD antibodies for western blotting. Indicated recombinant AtCYT6 proteins were loaded for size comparison. The elutions of the CoIPs displayed bands corresponding in size to the individual AtCYT6-NTDs or –CTD respectively.

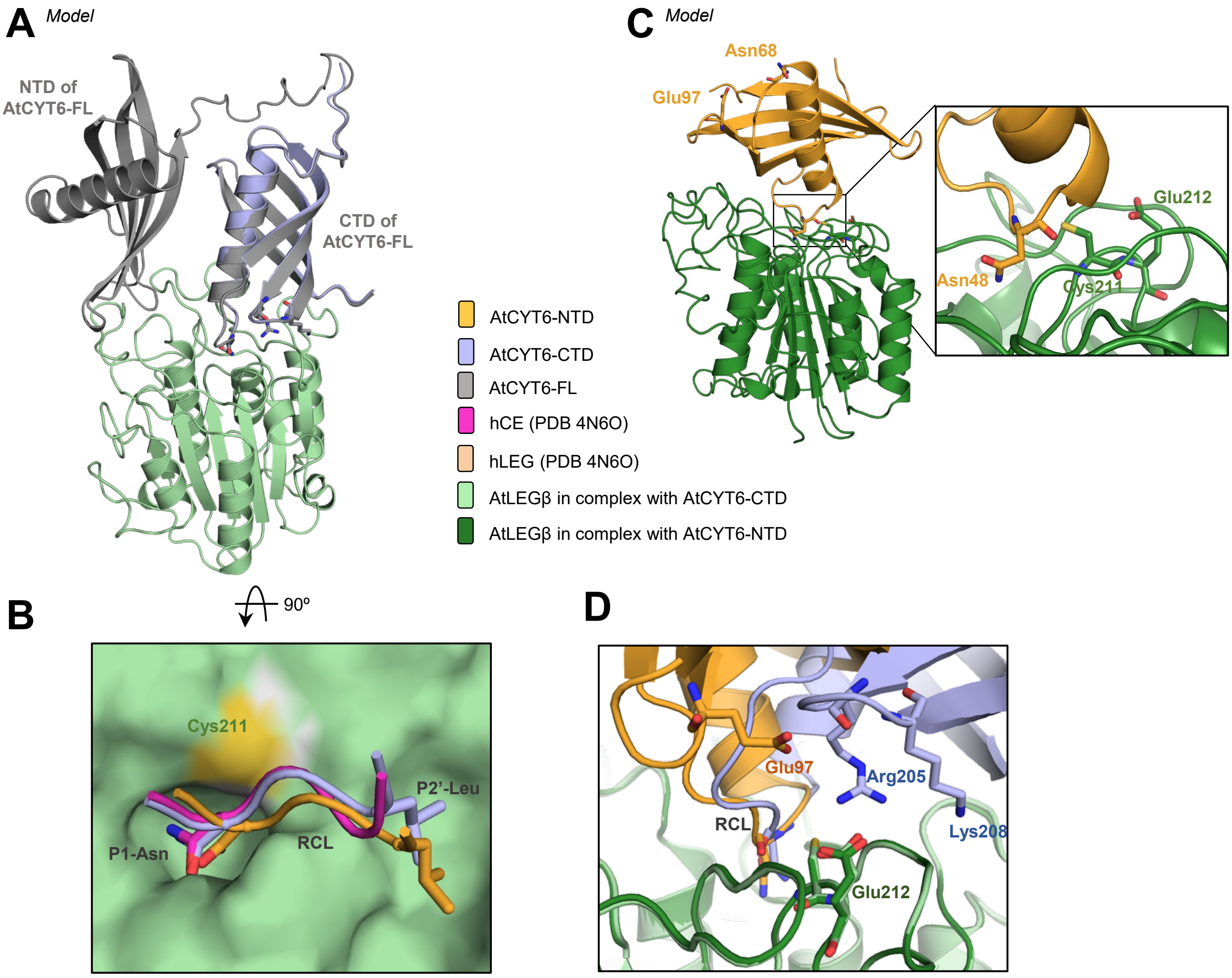


**Supplementary figure 4: Models of AtLEGβ in complex with AtCYT6-FL, AtCYT6-NTD or -CTD. (A)** Binding mode of AtCYT6-FL (grey) to AtLEGβ as predicted by AlphaFold Multimer. The CTD of the full-length protein binds to AtLEGβ (green) with the same orientation as AtCYT6-CTD (blue) and does not interact with AtCYT6-NTD. **(B)** Top-view on the RCL bound to the active site of AtLEGβ (green surface). Orange: AtCYT6-NTD, blue: AtCYT6-CTD, pink: hCE. **(C)** Binding mode of AtCYT6-NTD to AtLEGβ as originally predicted by AlphaFold Multimer. In the model, the AtCYT6-NTD (wheat) binds to AtLEGβ (green) with the Asn48 inserted into the active site, whereas Asn68 and Glu97 are positioned on the opposite side of the active site of AtLEGβ. **(D)** Zoom-in view on the LEL. In the model, residues Glu97 of AtCYT6-NTD (orange), and Arg205 and Lys208 of AtCYT6-CTD (blue) were in close proximity to Glu212 on AtLEGβ (green).

```
FcCYT1    MKLKS----QS----NSSPSFVVFFVLFFSF--------------IALSEMSTLGGIRDS 38
FaCYT1    MKLRS----QS----NSSPSFVFFFVLFFSF--------------IALSEMATLGGIRDS 38
AtCYT6    ---------------MMRSRFLLF-IVFFSL-SLFISSLIASDLGFCNEEMALVGGVGDV 43
OsCYT12   MRVAATTRPASSSAAAPLPLFLLL-AVAAAAAALFLVG--SASLA---MAGHVLGGAHDA 54
HvCYT4    MRVAAT-RPVSSA---PV-------ALLAALALLFLVG--SASLAIGTMASHVLGGKSEN 47
                                    :  :                       :**  :

FcCYT1    PAGSENSLETEALGRFAVDDHNQKQNGMLEFVRVVKAKEQVVSGTLHHLVVEAIDGGKKK 98
FaCYT1    PAGSENSLETEALGRFAVDDHNQKQNGMLEFVRVVKAKEQVVAGTLHHLVVEAIDGGKKK 98
AtCYT6    PA-NQNSGEVESLARFAVDEHNKKENALLEFARVVKAKEQVVAGTLHHLTLEILEAGQKK 102
OsCYT12   PS-AANSVETDALARFAVDEHNKRENALLEFVRVVEAKEQVVAGTLHHLTLEALEAGRKK 113
HvCYT4    PA-AANSLETDGLARFAVDEHNKRENALLEFVRVVEAKEQTVAGTLHHLTLEALEAGRKK 106
          *:   ** *.:.*.*****:**:::*.:***.***:****.*:******.:* ::.*:**

FcCYT1    LYEAKVWVKPWLNFKQVQEFKHAGDPETVSGTPSFTSSDLGVKQGGHAPGWQDVHPHDPQ 158
FaCYT1    LYEAKVWVKPWLNFKQVQEFKHAGEPETVSGTPSFTSSDLGVKQGGHAPGWQDVHPQDPQ 158
AtCYT6    LYEAKVWVKPWLNFKELQEFKPASDA------PAITSSDLGCKQGEHESGWREVPGDDPE 156
OsCYT12   VYEAKVWVKPWLDFKELQEFRNTGDA------TTFTNADLGAKKGGHEPGWRDVPVHDPV 167
HvCYT4    VYEAKVWVKPWLDSKELQEFRHTGDA------TSFTISDLGAKRGGHEPGWRDVPVHDPV 160
          :***********: *::***: :.:        ::* :*** *:* *  **::*  .**

FcCYT1    VQDAANHAVKSLQQKSNSLFPYELQEVVHAKAEVMEEHAKFNMLLKLKRGDKEEKYKVEV 218
FaCYT1    VQDAANHAVKSLQQKSNSLFPYELQEVVHAKSEVMEEHAKFNMLLKLKRGDKEEKYKVEV 218
AtCYT6    VKHVAEQAVKTIQQRSNSLFPYELLEVVHAKAEVTGEAAKYNMLLKLKRGEKEEKFKVEV 216
OsCYT12   VKDAADHAVKSIQQRSNSLFPYELLEIVRAKAEVVEDFAKFDILMKLKRGNKEEKFKAEV 227
HvCYT4    VKDAASHAVKSIQERSNSLFPYELIEIVRAKAEVVEDFAKFDIVMKLKRGTKEEKMKAEV 220
          *:..*.:***::*::********* *:*:**:**  : **:::::***** **** *.**

FcCYT1    HKNNEGAYNLNQMEVEH------ 235
FaCYT1    HKNNEGAYNLNQMEVEH------ 235
AtCYT6    HKNHEGALHLNHAEQHHD----- 234
OsCYT12   HKNLEGAFVLNQMQQEHDESSSQ 250
HvCYT4    HKNLEGAFVLNQMQPEHDESSSQ 243
          *** ***  **: : .*
```

**Supplementary figure 5: Sequence alignment of five type II phycys with demonstrated inhibitory activity against PLCPs and legumains.** Sequences were retrieved from Uniprot entries Q0JNR2 (OsCYT12, *Oryza sativa* cystatin 12), Q1ENE9 (HvCYT4, *Hordeum vulgare* cystatin 4), Q8H0X6 (AtCYT6, *A. thaliana* cystatin 6), Q4GZT8 (FaCYT1, *Fragaria ananassa* cystatin 1) and A0A2U9AEI9 (FcCYT1, *Fragaria chiloensis* cystatin 1). The P1-Asn residue on the RCL of the N- and C-terminal domains is indicated by solid and dashed black boxes, respectively. The conserved R-G-X-K double-basic motif on the LEL of the legumain-inhibitory CTDs is highlighted by a dashed grey box, and aspartate and glutamate residues on the LEL of the N-terminal domains by a solid grey box.

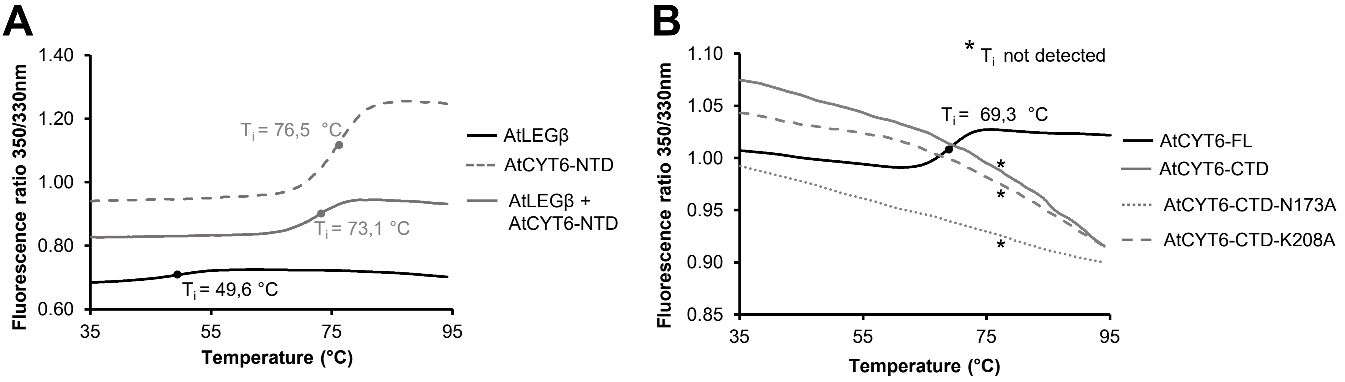


**Supplementary figure 6: Thermal stability of different AtCYT6 constructs.** The thermal stability of AtLEGβ and AtCYT6 constructs was measured by nanoDSF (differential scanning fluorimetry) measurements. Unfolding transitions (inflection point; $T_i$) are indicated by solid circles. **(A)** AtLEGβ was inhibited with MMTS and incubated at pH 6.5 in the absence (black) and presence of AtCYT6-NTD (dark grey). AtCYT6-NTD alone is shown as dashed grey line. **(B)** Denaturation curves obtained for AtCYT6-FL (solid black line), AtCYT6-CTD (grey solid line) AtCYT6-CTD-N173A (grey dotted line) and AtCYT6-CTD-K208A (grey dashed line) at pH 6.5 as controls for experiments depicted in Fig. 7.

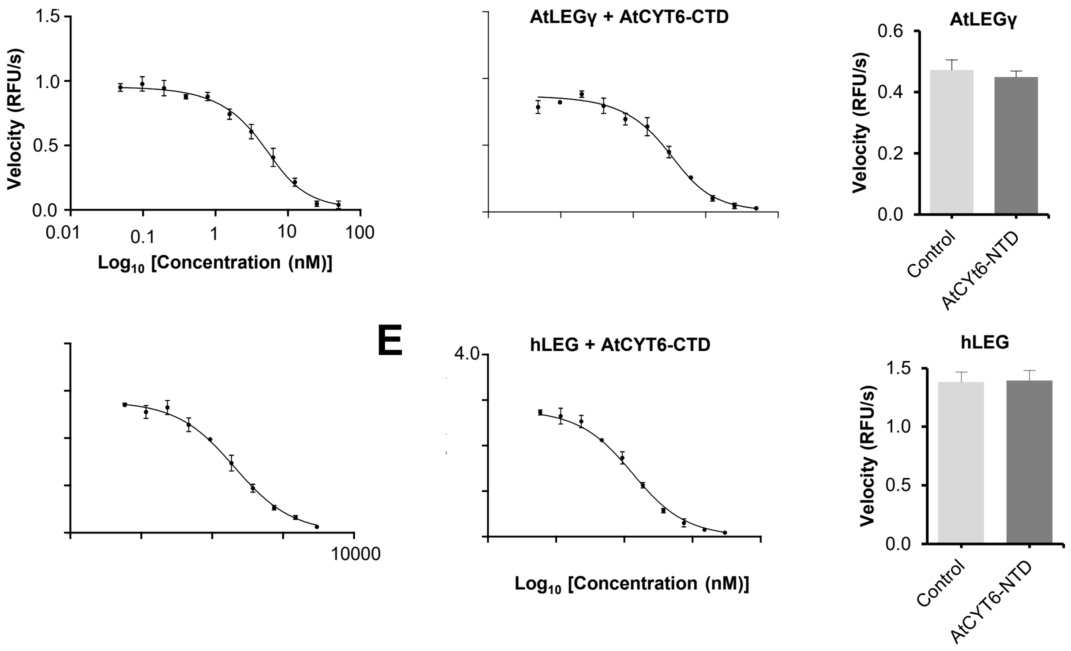


**Supplementary figure 7: Inhibition assays of AtCYT6 constructs against AtLEGγ and hLEG.** Enzymatic activity was measured at pH 5.5 as the increase in fluorescence after turnover of the AAN-AMC substrate. **(A)** and **(B)** The Ki of indicated AtCYT6 constructs towards AtLEGγ was determined using Morrison's equation. **(C)** Activity of AtLEGγ in the presence (dark grey) and absence (light grey) of AtCYT6-NTD. **(D)** and **(E)** The Ki of indicated AtCYT6 constructs towards hLEG was determined using Morrison's equation. **(F)** Activity of hLEG in the presence (dark grey) and absence (light grey) of AtCYT6-NTD.

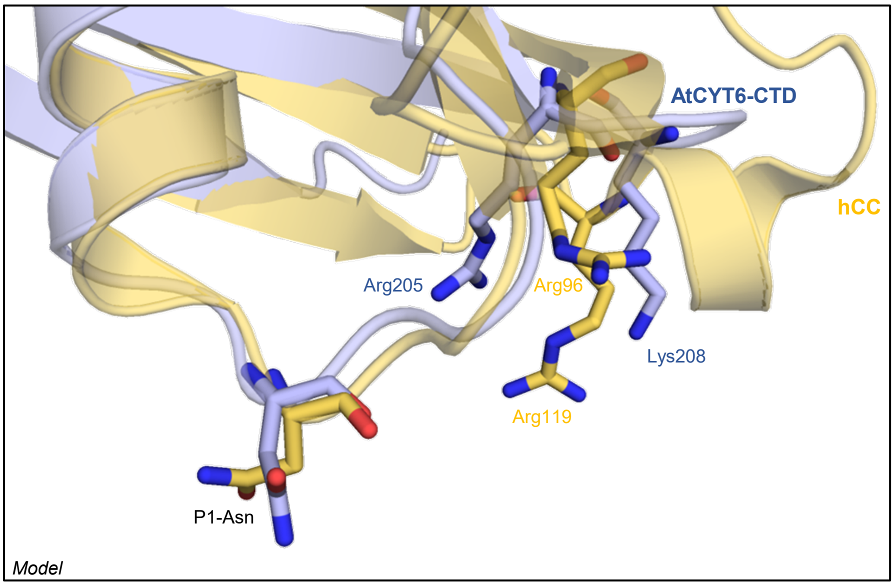


**Supplementary figure 8: Structure alignment of the modeled AtCYT6-CTD (blue) with human cystatin C (hCC, PDB 3GAX; yellow).** The P1-Asn residues and residues on the LEL are shown as sticks.

#### Supplementary Tables

**Supplementary Table 1: Mass spectrometry analysis (intact mass) of AtCYT6-derived constructs expressed in *E. coli*.**

| *Construct* (AtCYT6-) | *Theoretical mass (Da)* | *Experimental mass (Da)* | *Delta mass (Da)* | *Position* | *Sequence* | *Relative abundance (%)* |
|---|---|---|---|---|---|---|
| **FL** | - | no signal | - | - | - | - |
| **NTD** | 12611.46 | 12611.59 | -0.13 | 2-115 | (M)GSSH…PASD | 100 |
| **CTD** | 12465.169 | 12465.30 | -0.13 | 7-115 | (H)HHH…QHHD | 100 |
| **$CTD_{long}$** | 14299.01 | 14299.19 | -0.18 | 2-129 | (M)GSSH…HHD | 100 |

**Supplementary Table 2: Cleavage sites (P1 residues) of AtLEGβ within AtCYT6-NTD and AtCYT6-CTD identified by mass spectrometry analysis.**

| | **AtCYT6-NTD** | **AtCYT6-CTD** |
|---|---|---|
| **Cleavage sites** | Asn46<br>Asn68<br>Asn115 | Asn173 |
